# Sparsening and Decorrelation of Granule Cell Activity in the Dentate Gyrus by Noradrenaline

**DOI:** 10.1101/2025.07.08.662991

**Authors:** Iulia Glovaci, Koen Vervaeke, Hua Hu

## Abstract

The dentate gyrus in the hippocampus makes important contributions to the acquisition of episodic memories by transforming synaptic inputs from the entorhinal cortex into sparse and decorrelated activity patterns of its principal neurons, the granule cells. However, the underlying mechanism remains unclear. Using a combination of electrophysiological and optical recordings, together with optogenetic and pharmacological manipulations, we demonstrate that the release of noradrenaline plays a key role in this specialization via an enhancement of feedforward inhibition generated by cholecystokinin-expressing interneurons. By imposing coincidence detection with milliseconds temporal resolution onto granule cells, this enhancement of feedforward inhibition makes granule cell activity sparser and their firing patterns decorrelated. Since decorrelation contributes to efficient memory storage during auto-associative learning, these findings reveal a circuit mechanism by which an arousal signal facilitates memory formation in the hippocampus.

## Introduction

The dentate gyrus plays a key role in learning and memory. Theories of hippocampal computation and experimental evidence indicate that the dentate gyrus transforms synaptic inputs from the entorhinal cortex, which carry spatial and multisensory information, into decorrelated action potential (AP) patterns of its principal neurons (the granule cells)^1, 2, 3, 4, 5, 6, 7, 8^. This decorrelation operation at the encoding stage of associative learning is thought to be crucial for the efficient storage and accurate retrieval of episodic memories^6, 9^. Consequently, how granule cells transform afferent synaptic inputs to their APs is critically important for the formation of episodic memories in the hippocampus. This intricate process depends on a complex interplay between glutamatergic excitatory neurons and γ-aminobutyric acid (GABA)-releasing inhibitory interneurons^1, 2^, which is further controlled by several types of subcortical inputs that release neuromodulators.

In vivo electrophysiological and optical recordings from the dentate gyrus have demonstrated that AP discharge in granule cells is highly sparse and dynamically regulated by behavioral states. Specifically, the activity level of granule cells becomes sparser during arousal and attention^10, 11, 12^. Previous work suggests that this functional feature can contribute to memory formation by reducing the overlap between memory representations^6, 13, 14, 15^, but the mechanism underlying the behavioral state-dependent suppression of granule cell firing is not fully understood. A neuromodulatory circuit strongly engaged in these behavioral states is the noradrenergic system originating from the locus coeruleus^16^. Based on the dense innervation of the dentate gyrus by noradrenergic axons^17^, we hypothesize that the release of noradrenaline can sparsen and decorrelate granule cell activity.

Using a combination of electrophysiological and optical recordings, together with optogenetic and pharmacological manipulations, we demonstrate that noradrenaline suppresses the synaptic activation of granule cells by enhancing cholecystokinin-expressing interneuron (CCK^+^-IN)-mediated feedforward inhibition. This enhancement of feedforward inhibition, by imposing powerful coincidence detection onto granule cells, allows these principal neurons to be preferentially activated by highly synchronous excitatory synaptic inputs. Subsequent predictive computational modeling, supported by experimental evidence, indicates that noradrenaline harnesses this mechanism, allowing granule cells to recognize subtle differences in their excitatory inputs and generate decorrelated AP patterns.

## Results

### Noradrenaline inhibits the synaptic activation of granule cells

To induce noradrenaline release using an optogenetic approach, we virally expressed ChrimsonR fused to tdTomato in locus coeruleus noradrenergic neurons (**Fig. 1A**, **1B**)^18^. Four weeks post-surgery, strong expression of ChrimsonR-tdTomato was detected in axons reaching the dentate gyrus (**Fig. 1C**). By performing loose-patch recordings from tdTomato-labeled axons in the dentate gyrus in acute hippocampal slices (**Fig. 1D**), we found that suprathreshold photostimulation evoked tetrodotoxin-sensitive axonal APs, which were tightly coupled to individual light pulses (**Fig. 1E**–**F**, reliability for evoking an axonal AP = 100% under control conditions and 0% in 0.5 µM bath-applied tetrodotoxin, respectively; n = 8 axons from 6 mice). These subcellular recordings confirmed that the optical stimulation could activate noradrenergic axons with high fidelity and temporal precision.

**Figure 1.**
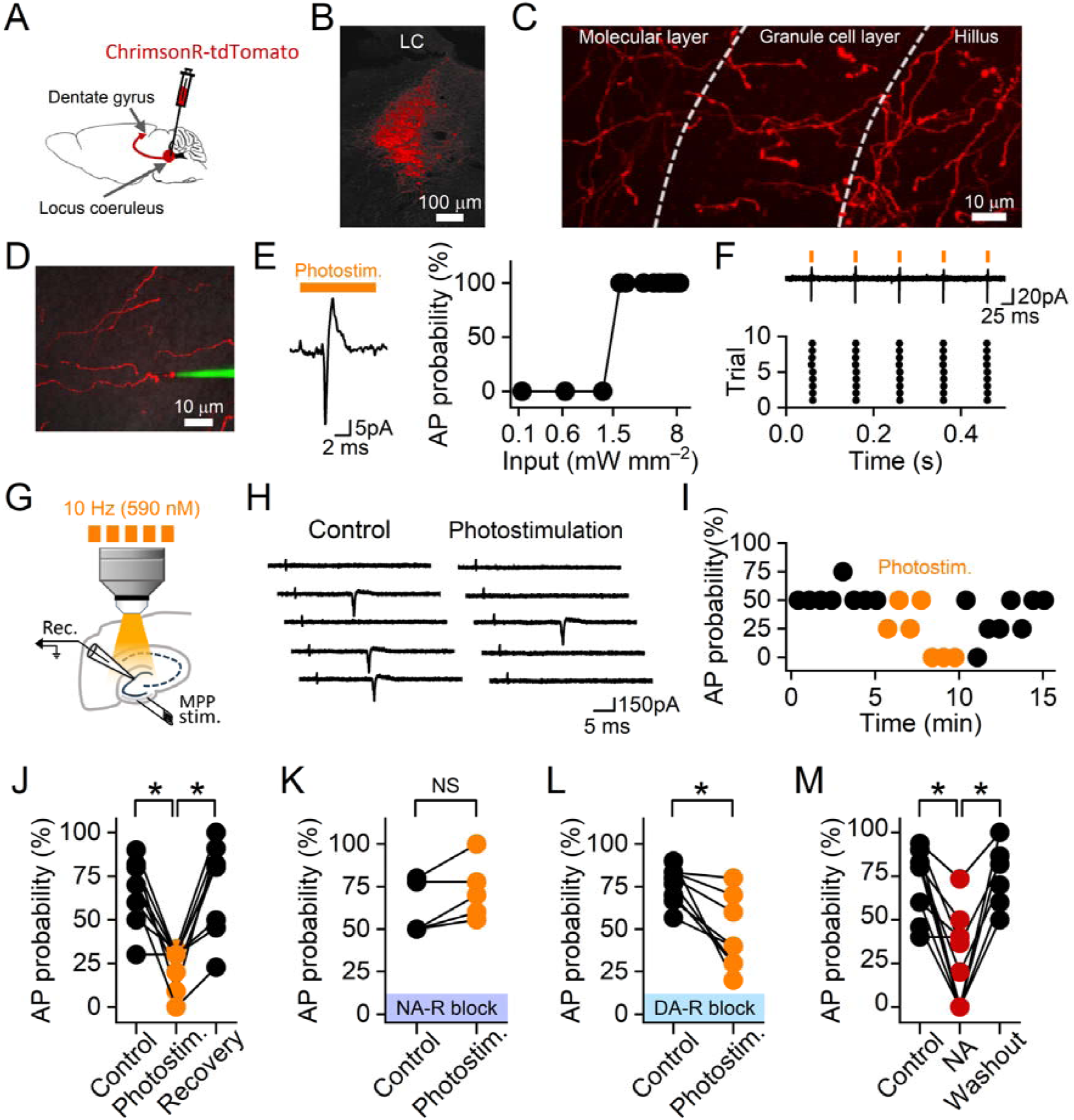
Noradrenaline inhibits the synaptic activation of individual granule cells. (**A**) Viral delivery of ChrimsonR-tdTomato to the locus coeruleus of TH-cre mice, illustrated schematically. (**B**) ChrimsonR-tdTomato expression in the locus coeruleus (LC). (**C**) A single confocal section showing ChrimsonR-tdTomato-expressing axons in an acute hippocampal slice prepared 4 weeks after the viral injection. (**D**) Loose-patch recording from a tdTomato-labelled axon with an electrode containing Alexa Fluor 488 (green). (**E**) Left, an axonal AP evoked by a 590-nm light pulse (5.2 mW mm^−2^, orange). Right, the probability of eliciting an axonal AP plotted against the light pulse intensity on a logarithmic scale. (**F**) Top, axonal APs evoked by a train of 10-Hz light pulses (5.2 mW mm^−2^, orange). Bottom, raster plot showing that each light pulse was coupled to an axonal AP during 9 consecutive trials. **D**–**F** are from the same experiment. (**G**) Experimental design for data presented in **H**–**L**, showing a cell-attached recording of granule cell APs evoked by electrically stimulating the medial perforant path (MPP stim.). Light pulses were used to optogenetically elicit the release of noradrenaline. (**H**) Responses of a granule cell to medial perforant path inputs under control conditions and following the optogenetic stimulation. Current traces from five consecutive trials were plotted for each condition. (**I**) The probability for medial perforant path inputs to activate the granule cell, plotted against time from the experiment in **H**, under control conditions (black) and when paired with the optogenetic activation of noradrenergic axons (orange). (**J**) Summary graph showing that the optogenetic stimulation of noradrenergic axons inhibited the synaptic activation of granule cells (n = 9 cells). (**K**–**L**) Similar to **J,** but results in **K** were acquired in bath-applied adrenergic receptor blockers (20 μM phentolamine and 50 μM propranolol, n = 6 cells). Experiments in **L** were performed in the presence of dopamine receptor blockers (50 μM SCH23390 and 50 μM sulpiride, n = 8 cells). (**M**) Summary graph indicates that bath-applied noradrenaline (NA) inhibited the synaptic activation of individual granule cells (n = 11 cells). In **J**–**M**, data points from the same experiment are connected by a line. * indicates P = 0.01, and NS indicates P = 0.07 (two-sided Wilcoxon signed-rank test).

To determine how the release of noradrenaline could modulate the activity of individual granule cells, we performed cell-attached recordings from these neurons. We activated granule cells by electrically stimulating medial perforant path axons that convey excitatory inputs from the medial entorhinal cortex to the dentate gyrus (**Fig. 1G**). We found that optogenetic activation of noradrenergic axons (five 10-ms-long light pulses at 10 Hz, paired with the electrical synaptic stimulation for 5 minutes) inhibited AP generation in granule cells evoked by the synaptic stimulation (**Fig. 1H**– **J**). The inhibition of granule cells, which was induced by the optogenetic activation of noradrenergic axons, was fully occluded by adrenergic receptor blockers but was resistant to dopaminergic receptor blockers (**Fig. 1K, L**), indicating that it depended on the activation of adrenergic receptors. Furthermore, the optogenetic activation of noradrenergic axons produced a similar degree of inhibition in the absence and presence of dopaminergic receptor blockers (P = 0.1, two-sided Wilcoxon rank sum test), suggesting that the inhibition of granule cells did not rely on the co-release of dopamine from noradrenergic axons^19^. In agreement, activating adrenergic receptors with bath-applied noradrenaline (10 µM) also inhibited the synaptic activation of granule cells (**Fig. 1M**).

How does noradrenaline influence granule cell activity at the population level? To answer this question, we performed extracellular recordings of population spikes that predominantly reflect the spiking activity of granule cells (**Fig. 2A**)^20^. Consistent with its effect on individual granule cells, optogenetic activation of noradrenergic axons reversibly reduced the size of the population spike (**Fig. 2B**–**E**), suggesting that noradrenaline inhibited the granule cell population. To confirm this observation in identified granule cells, we optically recorded their activity following the viral expression of GCaMP8s in these neurons (**Fig. 2F**). To separate Ca^2+^ events associated with APs from subthreshold Ca^2+^ signals^21^, we established a criterion by correlating simultaneously recorded GCaMP8s signals with electrophysiologically identified APs (**Supplementary Fig. 1**)^22^. Based on this criterion, we demonstrated that the occurrence of suprathreshold Ca^2+^ signals following the stimulation of perforant path axons became less frequent in noradrenaline (**Fig. 2G**–**I**).

**Figure 2.**
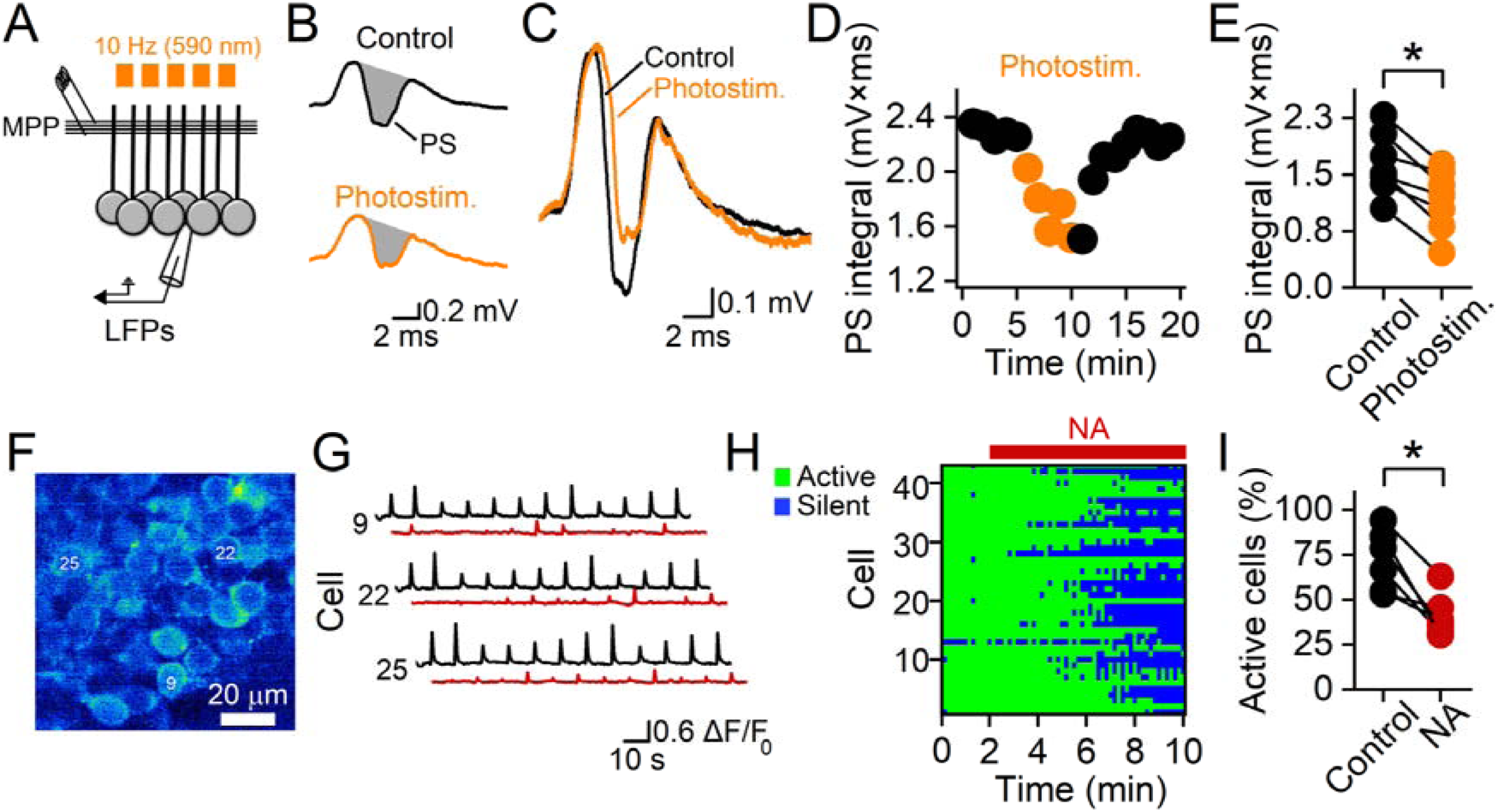
Noradrenaline inhibits the synaptic activation of the granule cell population. (**A**) Schematic diagram showing the experimental design for the recording of local field potentials (LFPs) to measure granule cell population activity evoked by electrically stimulating the medial perforant path (MPP). Optogenetic activation of noradrenergic axons was elicited with a train of light pulses (orange). (**B**) Local field potentials in the granule cell layer under control conditions (black) and following the optogenetic activation of noradrenergic axons (orange). The population spike (PS) is highlighted in grey. (**C**) Superimposition of local field potential traces in **B**. (**D**) Integral of the population spike plotted against time for the experiment illustrated in **B**–**C**. Orange circles represent data points obtained when the stimulation of perforant path axons was paired with optogenetic activation of noradrenergic axons. (**E**) Summary graph showing that optogenetic activation of noradrenergic axons inhibited the population spike (n = 7 experiments). (**F**) Confocal image showing the expression of GCaMP8s in granule cells in an acute hippocampal slice. Numbers indicate neurons selected for the illustration in **G**. (**G**) GCaMP8s signals evoked by stimulating medial perforant path axons in three representative granule cells. Black traces, GCaMP8s signals under control conditions, red traces, GCaMP8s signals during the bath application of noradrenaline. (**H**) Synaptic activation of 42 GCaMP8s-expressing granule cells plotted as a function of time in a binary fashion. Each green symbol indicates that the synaptic stimulation successfully activated a granule cell, whereas each blue symbol indicates a failure for the synaptic stimulation to activate a specific granule cell. Active granule cells were identified based on the amplitude of the Ca^2+^ signal evoked by the synaptic stimulation, using the criterion established in **Supplementary Fig. 1.** Data in **F**–**H** are from the same experiment. (**I**) Summary graph showing that bath-applied noradrenaline reduced the fraction of GCaMP8s-expressing granule cells that were activated by the synaptic stimulation (n = 6 experiments). Granule cells that did not display any suprathreshold Ca^2+^ activities under control conditions were excluded from this analysis. In **E** and **I**, data points from the same experiment are connected by a line. * indicates that P = 0.02 (two-sided Wilcoxon signed-rank test).

Collectively, these experiments provide complementary evidence to indicate that noradrenaline suppresses the activity of granule cells at both single-cell and population levels.

## Noradrenaline facilitates feedforward inhibition

We sought to identify the mechanism by which noradrenaline inhibits granule cells, and we obtained three lines of evidence to implicate synaptic inhibition. First, noradrenaline had no significant effect on the resting membrane potential or the input resistance of granule cells (resting membrane potential = 79.6 ± 1.8 mV under control conditions and 81.0 ± 1.9 mV in noradrenaline, n = 7, P = 0.24; input resistance = 242.2 ± 38.1 MΩ under control conditions and 227.5 ± 31.3 MΩ in noradrenaline, n = 7, P = 0.24, two-sided Wilcoxon signed rank-test). Furthermore, it did not modulate the voltage threshold of granule cell APs (–40.6 ± 1.9 mV under control conditions and –41.7 ± 2.1 mV in noradrenaline, n = 6, P = 0.08, two-sided Wilcoxon signed rank-test). These results suggest that noradrenaline does not inhibit granule cells by downregulating their intrinsic excitability. Second, bath application of noradrenaline did not affect excitatory postsynaptic currents (EPSCs) in granule cells evoked by stimulating medial perforant path axons (EPSC charge = 3.7 ± 0.5 pC under control conditions and 3.2 ± 0.6 pC in noradrenaline, n = 7, P = 0.13, two-sided Wilcoxon signed rank-test), indicating that noradrenaline did not inhibit granule cells by diminishing their excitatory drive. Third, preincubation of slices with a GABA_A_ receptor antagonist (SR95531, 4 µM) prevented the inhibition of granule cells by noradrenaline (**Fig. 3A**–**C**). Collectively, these experiments indicate that synaptic inhibition is essential for the suppression of granule cell activity by noradrenaline.

**Figure 3.**
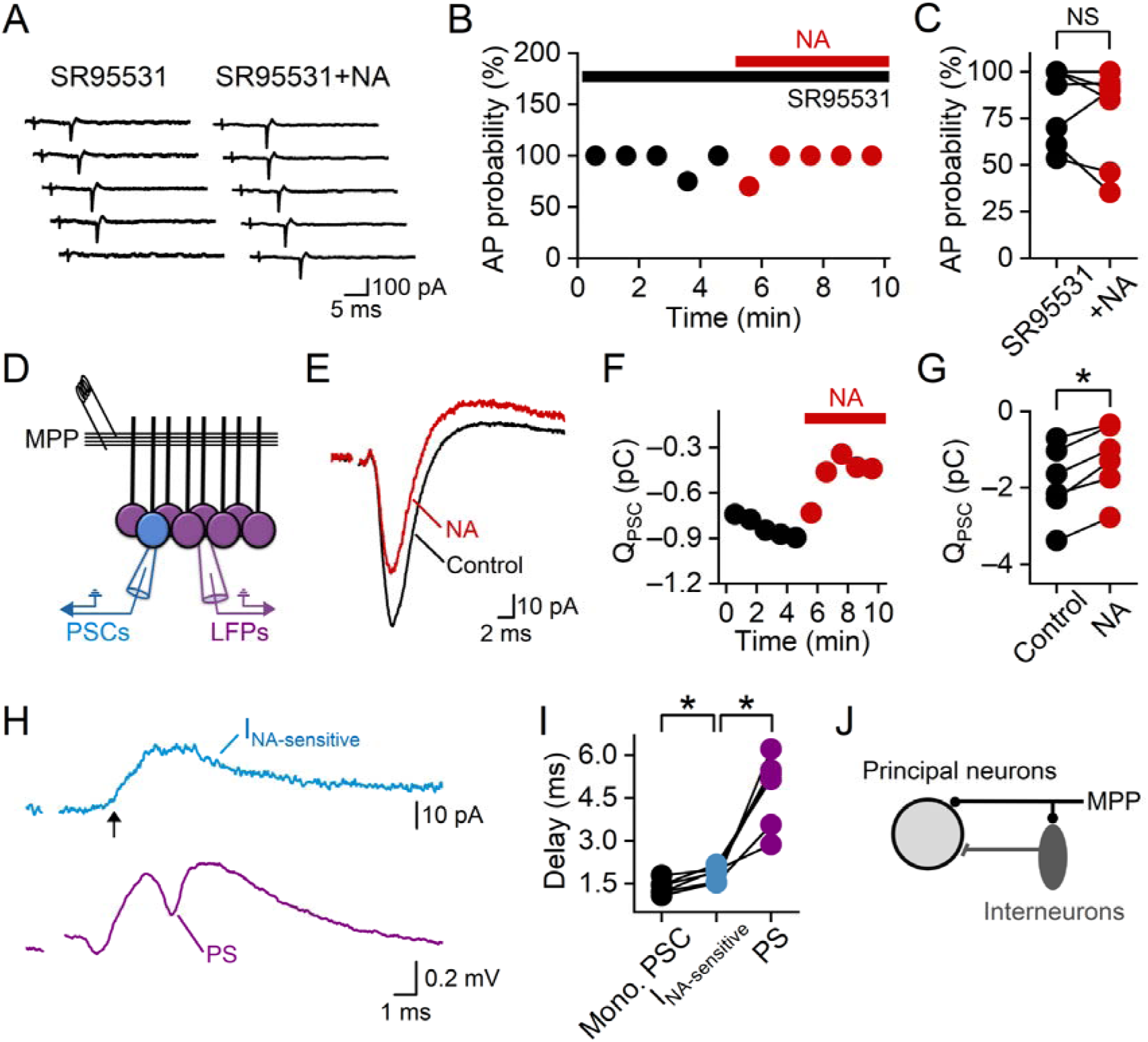
The suppression of granule cell activity by noradrenaline depends on synaptic inhibition. (**A**) Cell-attached recording showing that noradrenaline did not block the synaptic activation of the granule cell in the presence of bath-applied SR95531. Current traces from five consecutive trials were plotted for each condition. (**B**) The probability for medial perforant path inputs to evoke an AP in the granule cell plotted against time from the experiment in **A**. (**C**) Summary graph showing that SR95531 prevented the inhibition of granule cell activity by bath-applied noradrenaline. NS indicates that P = 0.29 (n = 6 cells, two-sided Wilcoxon signed-rank test). (**D**) Schematic diagram of the experimental design for the data presented in **E**–**I**, illustrating a simultaneous recording of postsynaptic currents (PSCs) in a granule cell and local field potentials (LFPs) in the granule cell layer evoked by stimulating the medial perforant path (MPP). (E) Postsynaptic currents in a granule cell under control conditions (black) and during the bath application of noradrenaline (red). (**F**) Net charge of postsynaptic currents plotted against time from the experiment in **E**. (**G**) Summary graph showing the effect of bath-applied noradrenaline on the net charge of postsynaptic currents. * indicates that P = 0.03 (n = 6 cells, two-sided Wilcoxon signed-rank test). (**H**) Noradrenaline-sensitive synaptic current (blue) in a granule cell and local field potential (purple) in the granule cell layer evoked by medial perforant path inputs. The arrow indicates the onset of the noradrenaline-sensitive current, which preceded the simultaneously evoked population spike (PS). The noradrenaline-sensitive current was obtained by subtracting the postsynaptic current under control conditions from that during the application of noradrenaline. (**I**) Summary graph comparing the onset of the noradrenergic-sensitive synaptic currents with those of the monosynaptic currents (Mono. PSC) and the population spikes, using the timing of the synaptic stimulation as a temporal reference. * indicates that P = 0.03 (n = 6 experiments, two-sided Wilcoxon signed-rank test). (**J**) Schematic diagram of the circuit motif underlying feedforward inhibition. In **C** and **G**, data points from the same experiment are connected by a line.

We subsequently determined the circuit mechanism by which noradrenaline could enhance synaptic inhibition in granule cells (**Fig. 3D**). Following the stimulation of medial perforant path axons, postsynaptic currents (PSCs) in granule cells consisted of a short-latency monosynaptic EPSC that partially overlaps with a delayed disynaptic inhibitory postsynaptic current (IPSC)^22^. We found that noradrenaline enhanced the disynaptic IPSC. This result is supported by two sets of experiments. First, the reversal potential of the noradrenaline-sensitive synaptic current (–72.0 ± 2.6 mV after the liquid junction potential compensation, n = 5), which was obtained by subtracting the postsynaptic current under control conditions from that in noradrenaline (**Fig. 3E**–**G**), was near the Cl^−^ reversal potential predicted by the Nernst equation (–71 mV), suggesting that it was generated by the activity of GABA_A_ receptors. Second, the noradrenaline-sensitive synaptic current occurred after the monosynaptic response but preceded the population spike (**Fig. 3H**–**I**). This distinct temporal feature implies that this current is a disynaptic response generated by interneurons, which fire APs earlier than granule cells in response to afferent inputs^23^. In agreement, noradrenaline was ineffective in modulating monosynaptic IPSCs evoked by directly stimulating interneuron axons (IPSC charge = 3.0 ± 0.8 pC under control conditions and 3.4 ± 1.0 pC in noradrenaline, n = 7, P =0.18, two-sided Wilcoxon signed rank-test). Collectively, these results indicate that the inhibition of granule cells by noradrenaline requires the activation of a feedforward inhibitory circuit motif in the dentate gyrus (**Fig. 3J**)^23, 24^.

### The noradrenergic inhibition of granule cells is independent of parvalbumin-expressing interneurons

We subsequently identified the interneuron subtype that mediates the noradrenergic enhancement of feedforward inhibition. The dentate gyrus neuronal network includes multiple subtypes of inhibitory interneurons^25, 26^. Because the noradrenergic inhibition of granule cells depends on the activation of a feedforward inhibitory circuit motif by medial perforant path axons, we focused on interneuron subtypes that are innervated by medial perforant path axons^25^. Interneurons that are specialized in feedback inhibition in the dentate gyrus, for example, somatostatin-expressing interneurons, are not included in subsequent analyses because their dendrites do not contact medial perforant path axons^25^.

We first analyzed parvalbumin-expressing interneurons (PV^+^-INs, **Supplementary Fig. 2A–C**), which make significant contributions to feedforward inhibition in the dentate gyrus^27^. Unexpectedly, we found that noradrenaline inhibited the activation of PV^+^-INs by synaptic inputs from medial perforant path axons (**Supplementary Fig. 2D–F**). This experiment argues against the idea that noradrenaline inhibits granule cells by facilitating the synaptic recruitment of PV^+^-INs. Furthermore, paired recordings from monosynaptically connected PV^+^-INs and granule cells indicated that noradrenaline did not enhance the release of GABA from PV^+^-INs (**Supplementary Fig. 2G–I**). Collectively, these experiments suggest that the inhibition of granule cells by noradrenaline is independent of PV^+^-INs. In agreement, neither chemogenetic (**Supplementary Fig. 2J–L)** nor pharmacological inhibition (DAMGO, 500 nM, **Supplementary Fig. 2M**) of PV^+^-INs prevented the suppression of granule cell activity by noradrenaline^28^. Furthermore, these pharmacological experiments also suggest that the noradrenergic suppression of granule cell activity does not require feedforward inhibition mediated by neurogliaform interneurons due to the inhibition of this interneuron subtype by DAMGO^29, 30, 31^.

### The noradrenergic inhibition of granule cells requires feedforward inhibition mediated by CCK^+^-INs

Based on the distribution of CCK^+^-IN apical dendrites in the molecular layer (**Fig. 4A**– **C**)^25^, we hypothesized that they contribute to feedforward inhibition in the dentate gyrus. Given that low levels of CCK expression have been detected in glutamatergic excitatory neurons and diverse types of GABAergic interneurons^32, 33, 34, 35^, we used an intersectional approach followed by rigorous electrophysiological and morphological classification to specifically target dentate gyrus CCK^+^-INs^36^. We found that APs in CCK^+^-INs always preceded granule cell spikes following the stimulation of medial perforant path axons (**Fig. 4D**–**E**). This temporal sequence indicated that these interneuron APs were not driven by feedback excitatory inputs from granule cells. Consistent with this idea, CCK^+^-IN-mediated disynaptic IPSCs, isolated with a type 1 cannabinoid receptor agonist (ACEA, 500 nM; **Fig. 4F**–**H**)^28, 37, 38^, preceded and inhibited granule cell APs (**Fig. 4I**–**M**). Collectively, these results indicated that dentate CCK^+^-INs could be activated by monosynaptic excitatory inputs from the medial entorhinal cortex and generate inhibition to oppose the excitatory drive. Such an operation is independent of granule cell activity and can only be performed by feedforward inhibitory circuits^23^.

**Figure 4.**
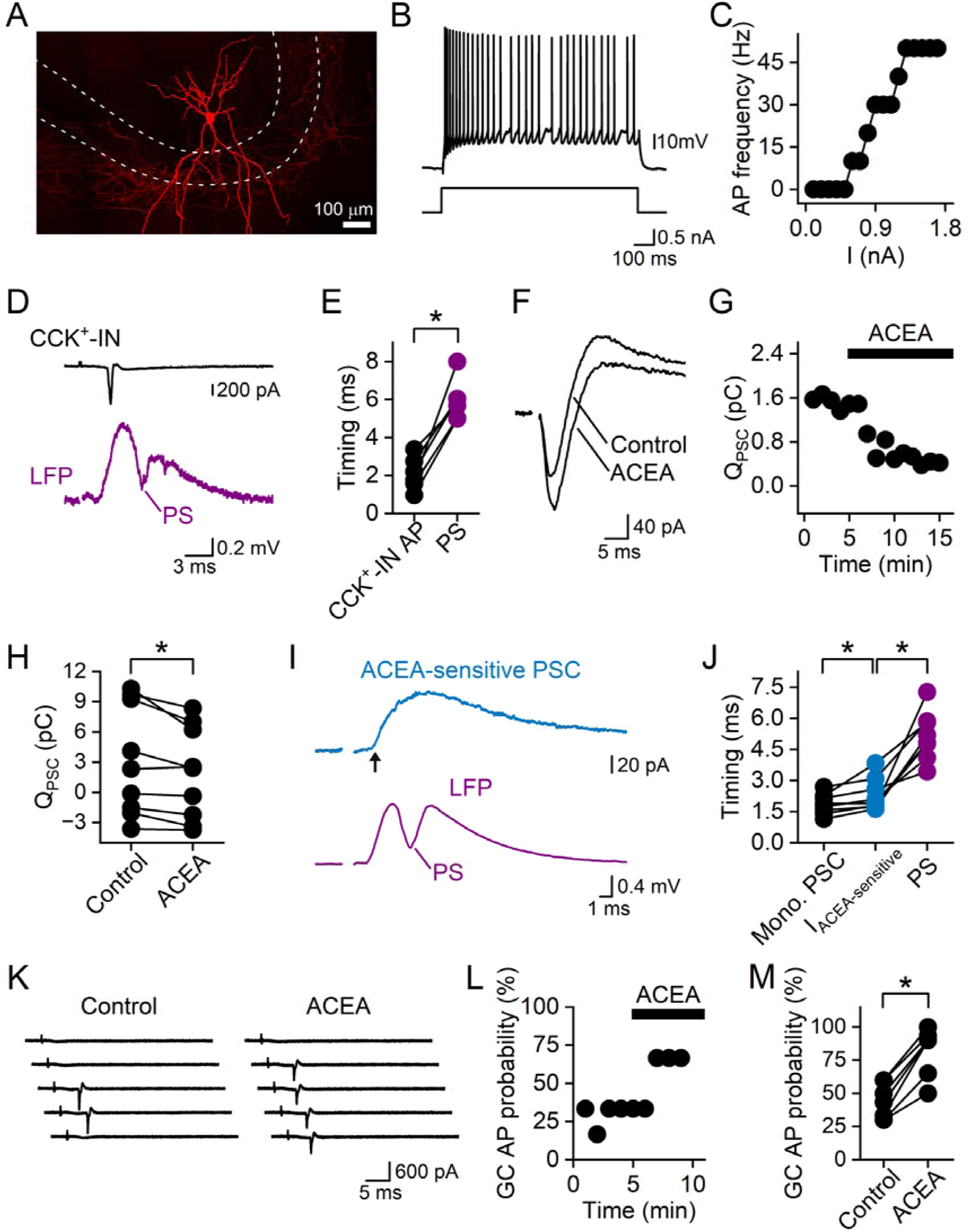
CCK^+^-INs contribute to feedforward inhibition in the dentate gyrus. (**A**) Morphology of a CCK^+^-IN in the dentate gyrus. White dashed lines indicate the borders of the granule cell layer. (**B**) APs (top) in a CCK^+^-IN evoked by the somatic injection of a 1-s depolarizing current pulse (bottom). (**C**) Steady-state AP frequency plotted against the amplitude of the current pulse from the experiment in **B**. (**D**) Simultaneous cell-attached recording of an AP in a CCK^+^-IN (black) and extracellular recording of local field potentials (LFPs, purple) in the granule cell layer evoked by stimulating the medial perforant path. (**E**) Summary graph comparing the timing of CCK^+^-IN APs with that of the population spike (PS), relative to the timing of the synaptic stimulation. * indicates that P = 0.03 (n = 6 experiments, two-sided Wilcoxon signed-rank test). (**F**) Superimposition of the postsynaptic currents in a granule cell under control conditions and during the application of a CB1R agonist (ACEA). (**G**) Net charge of postsynaptic currents plotted against time for the experiment in **F**. (**H**) Summary graph comparing the charge of the postsynaptic currents under control conditions with that in ACEA. * indicates that P = 0.02 (n = 7 cells, two-sided Wilcoxon signed-rank test). (**I**) ACEA-sensitive synaptic current (blue) and simultaneously evoked population spike (purple) in the granule cell layer, demonstrating that the onset of the ACEA-sensitive current (arrow) precedes the population spike. (**J**) Summary graph showing the onset of ACEA-sensitive postsynaptic currents (blue), the onset of monosynaptic currents (black), and the timing of the peak of population spikes (purple), relative to the synaptic stimulation artifact. * indicates that P = 0.01 (n = 8 experiments, two-sided Wilcoxon signed-rank test). (**K**) Cell-attached recording from a granule cell, showing the response to medial perforant path inputs under control conditions and during the bath application of ACEA. Five consecutive trials (10-s interval) were plotted for each condition. (**L**) The probability of medial perforant path inputs to elicit an AP in the granule cell in **K,** plotted against time. (**M**) Summary graph demonstrating that ACEA increased the probability for medial perforant path inputs to evoke an AP in granule cells. * indicates that P = 0.02 (n = 7 cells, two-sided Wilcoxon signed-rank test). In **H**, **J**, and **M**, data points from the same experiment are connected by a line.

We subsequently sought to determine whether CCK^+^-INs could be modulated by noradrenaline. We found that noradrenaline produced a significant depolarization of the resting membrane potential of CCK^+^-INs (**Fig. 5A**–**B**, –62.0 ± 0.9 mV under control conditions and –52.8 ± 1.4 mV in noradrenaline, P = 0.01, n = 13 cells, two-sided Wilcoxon signed-rank test). This depolarization is expected to facilitate the synaptic activation of CCK^+^-INs. In close agreement, we found that noradrenaline increased the probability for medial perforant path inputs to activate CCK^+^-INs (**Fig. 5C**–**E**). However, noradrenaline had no significant effect on the unitary IPSC (uIPSC) generated by CCK^+^-INs (**Fig. 5F**–**H**). Thus, noradrenaline produced an opposite effect on the synaptic recruitment of CCK^+^-INs and granule cells. This dichotomy suggested that noradrenaline might inhibit granule cells by facilitating synaptic activation of CCK^+^-INs. Based on this idea, we predicted that pharmacological inhibition of CCK^+^-INs would compromise the noradrenergic inhibition of granule cells. Indeed, ACEA prevented the inhibition of granule cells by optogenetic activation of noradrenergic axons and by bath-applied noradrenaline (**Fig. 5I**–**L**).

**Figure 5.**
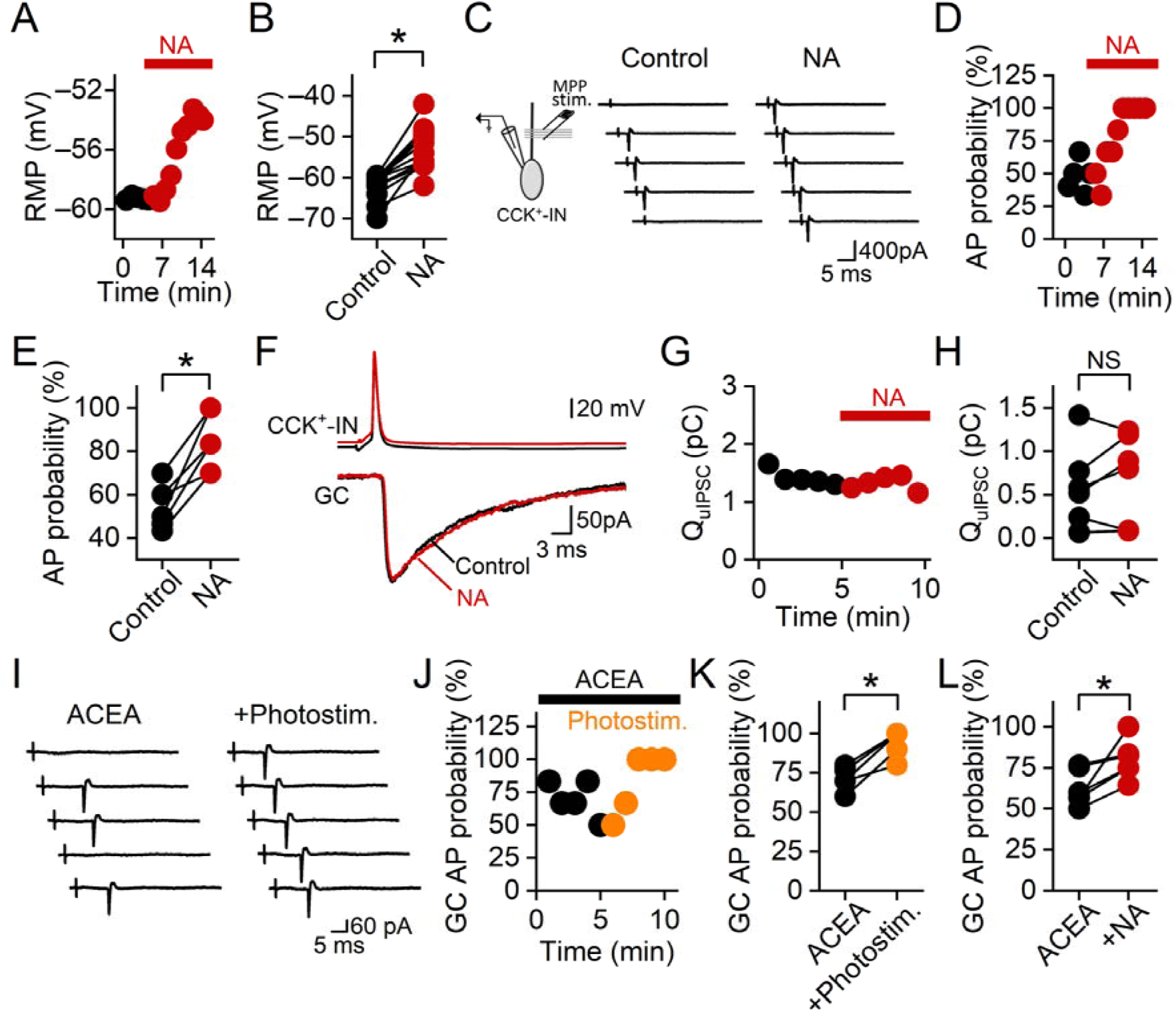
The inhibition of granule cells by noradrenaline is dependent on CCK^+^-INs. (**A**) Resting membrane potential value of a CCK^+^-IN plotted against time before and during the bath application of noradrenaline. (**B**) Summary graph comparing the resting membrane potential of CCK^+^-INs under control conditions with that in noradrenaline. ** indicates that P = 0.01 (n = 13 cells, two-sided Wilcoxon signed-rank test). (**C**) Cell-attached recording of APs in a CCK^+^-IN evoked by stimulating medial perforant path axons, before and during bath application of noradrenaline. (**D**) The probability of the synaptic stimulation to evoke an AP in the CCK^+^-IN plotted against time from the experiment in **C**. (**E**) Summary graph showing that bath-applied noradrenaline increased the probability of activating CCK^+^-INs by the synaptic stimulation. * indicates that P = 0.03 (n = 6 cells, two-sided Wilcoxon signed-rank test). (**F**) Top, APs in a CCK^+^-IN under control conditions (black) and during bath application of noradrenaline (red). Bottom, corresponding uIPSCs in a granule cell. (**G**) uIPSC charge plotted against time from the experiment in **F**. (**H**) Summary graph comparing the charge of CCK^+^-IN-mediated uIPSCs in granule cells under control conditions with that in noradrenaline. NS indicates P = 0.75 (n = 6 experiments, two-sided Wilcoxon signed-rank test). (**I**) Cell-attached recording of APs in a granule cell evoked by stimulating medial perforant path axons under control conditions and following the optogenetic activation of noradrenergic axons. ACEA was applied to the bath throughout the experiment to inhibit the release of GABA from CCK^+^-INs. (**J**) The probability for the synaptic stimulation to evoke an AP in the granule cell plotted against time from the experiment in **I**. (**K**) Summary graph showing that ACEA prevents the suppression of granule cell activity by optogenetic stimulation of noradrenergic axons. * indicates that P = 0.03 (n = 6 cells, two-sided Wilcoxon signed-rank test). Black circles, data points in ACEA; orange circles, data points following the optogenetic stimulation of noradrenergic axons in ACEA. (**L**) Summary graph showing that ACEA blocks the suppression of granule cell activity by bath-applied noradrenaline. * indicates that P = 0.03 (n = 6 cells, two-sided Wilcoxon signed-rank test). In summary graphs (**B**, **E**, **H**, **K**, and **L**), data points from the same experiment are connected by a line.

In the dentate gyrus, CB1 receptors are also expressed by mossy cells^39^. We believe that the effect of ACEA in our experiments is independent of mossy cells for three reasons. First, our synaptic stimulation electrode was positioned in the middle third of the molecular layer, which is spatially segregated from mossy cell axons and dendrites^39^. Thus, the probability that we directly stimulated mossy cells is extremely low. Second, ACEA-sensitive synaptic currents in our experiments display a short delay following the activation of the medial perforant path axons (**Fig. 4J**), arguing against that they were generated by the slower recurrent feedback synaptic response of mossy cells. Third, the recurrent feedback synaptic response of mossy cells, which depends on excitatory inputs from granule cells^39^, cannot precede granule cell APs in our experiments. Collectively, these three lines of evidence argue against a potential contribution from mossy cells to the effect of ACEA in our experiments.

### Noradrenaline inhibits granule cell firing evoked by repetitive stimulation of medial perforant path axons

In vivo electrophysiological recordings indicate that granule cells receive trains of EPSCs at the high gamma-frequency band (mean frequency = 115 Hz), which are nested inside theta-frequency oscillations, during spatial exploration^40, 41^. To determine whether noradrenaline can suppress granule cell APs evoked by excitatory synaptic inputs with this temporal feature, we delivered short trains of 115-Hz electrical stimulation to the medial perforant path, with successive trains separated by a 125-ms interval (**Supplementary Fig. 3A**). Consistent with previous studies^42, 43^, the probability for individual stimuli to activate the granule cell decreased in an activity-dependent manner during the repetitive synaptic stimulation, making granule cells preferentially activated by the early stimuli, and bath-applied noradrenaline inhibited the APs evoked by the stimulation protocol (**Supplementary Fig. 3B–D**). To identify the underlying synaptic mechanism, we performed analyses of PSCs evoked by this stimulation protocol (**Supplementary Fig. 3E–G**). We found that noradrenergic-sensitive IPSCs displayed a fast onset (1.90 ± 0.04 ms, n = 6 experiments) and fully dissipated before the next train of synaptic stimulation (current amplitude before the next train = 1.6 ± 1.7 pA, n = 6 experiments), suggesting a strong contribution from feedforward inhibition.

### Noradrenaline imposes coincidence detection onto granule cells

Because synaptic activation of individual granule cells requires spatial and temporal summation of a large number of excitatory synaptic inputs^44^, we sought to determine whether the modulation of feedforward inhibition by noradrenaline could influence the integrative properties of individual granule cells.

In response to afferent inputs, disynaptic feedforward inhibition arrives with a longer delay when compared to the monosynaptic excitation (**Fig. 6A**)^23^. This delayed arrival of synaptic inhibition creates a “window of opportunity” for synaptic excitation to activate granule cells, given that within this window synaptic inhibition is not sufficiently strong to prevent the depolarization generated by the excitation^23, 45^. Because noradrenaline enhances feedforward inhibition in the dentate gyrus, we predict that the modulation will make this window narrower. A narrow window allows granule cells to be preferentially activated by synchronous excitatory inputs.

**Figure 6.**
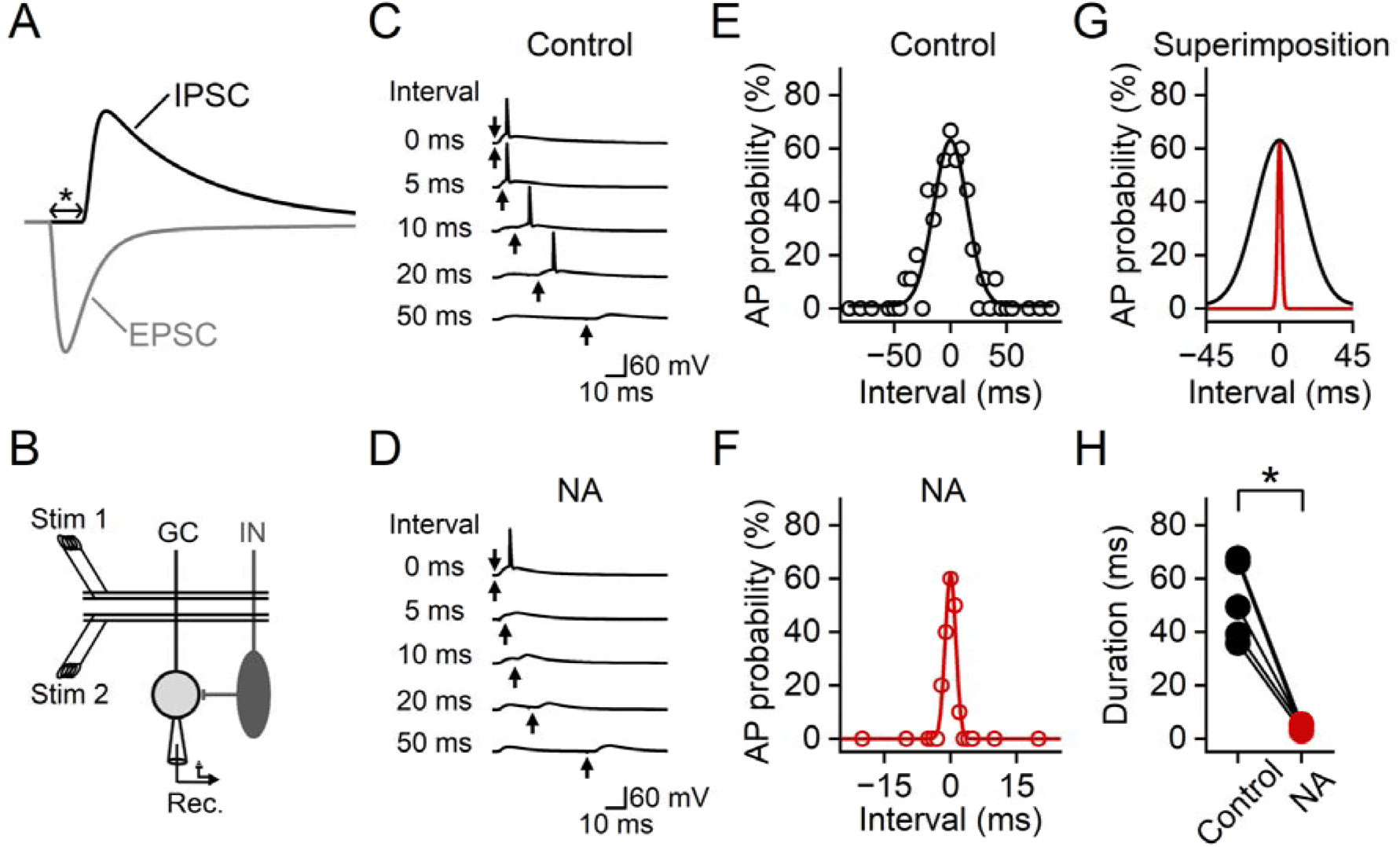
Noradrenaline imposes coincidence detection onto granule cells. (**A**) Schematic diagram illustrating the delay (indicated by the asterisk) between the onset of the monosynaptic EPSC and that of the disynaptic feedforward IPSC. (**B**) Schematic diagram of the experimental design for the data presented in **C**–**H**, illustrating whole-cell recording from a granule cell to measure the integration of two independent excitatory inputs evoked by stimulating medial perforant path axons. The independence was confirmed by demonstrating that stimulating one axonal pathway did not induce short-term plasticity on the EPSC generated by the activation of the other axonal pathway 60 ms later. (**C**) Membrane potential of a granule cell evoked by the two synaptic inputs. Arrows indicate the timing of these two inputs, and numbers on the left indicate the interval between them, which was determined by using the timing of the top arrow as the temporal reference. Positive intervals indicate that the input represented by the bottom arrow followed the one represented by the top arrow, and negative intervals in **E**–**G** indicate the reverse temporal order. (**D**) Similar to **C**, but in bath-applied noradrenaline. To compensate for the reduction of AP probability in noradrenaline, the synaptic stimulation intensity was increased to ensure that simultaneous activation of both axonal pathways evoked a postsynaptic AP with a probability matching that under control conditions (54.4 ± 3.9% under control conditions and 48.3 ± 3.1% in noradrenaline, P = 0.22, n = 6 cells, two-sided Wilcoxon signed-rank test). (**E**–**F**) The probability for the summation of the two synaptic inputs to evoke a postsynaptic AP plotted against the interval between them under control conditions (**E**) and in noradrenaline (**F**). Lines represent Gaussian functions fit to the data points. (**G**) Superimposition of Gaussian functions in **E** (black) and **F** (red). **C**–**G** are from the same experiment. (**H**) Summary plot indicating that the duration of the granule cell integration window in noradrenaline (4.0 ± 0.4 ms, red circles) is shorter than that under control conditions (51.1 ± 5.9 ms, black circles, * indicates P = 0.03, n = 6 cells, two-sided Wilcoxon signed-rank test). The duration was quantified from the width of the Gaussian function at its half-maximal amplitude in each experiment. Data points from the same experiment are connected by a line.

To test this idea, we measured the integration window during which the summation of two independent subthreshold excitatory inputs could activate single granule cells (**Fig. 6B**). By systematically changing the interval between these two inputs, we found that noradrenaline narrowed the granule cell integration window by an order of magnitude. Under control conditions, granule cells could be activated by excitatory synaptic inputs that were separated by tens of milliseconds (**Fig. 6C** and **E**). In bath-applied noradrenaline, these inputs were no longer effective in activating granule cells when the interval between them was longer than 5 ms (**Fig. 6D** and **F**). This sharp contrast (**Fig. 6E**–**H**) confirms that noradrenaline imposes a narrow integration window on granule cells, providing them with the ability to discriminate between highly synchronous inputs and temporally dispersed asynchronous inputs.

### Facilitation of feedforward inhibition enhances decorrelation

Theories of hippocampal computation propose that a key function of the dentate gyrus is to convert excitatory inputs from the entorhinal cortex into decorrelated granule cell AP patterns^1, 3, 6^. We hypothesized that noradrenaline could facilitate this specialization, given that it enhances the ability of granule cells to discriminate inputs with different temporal structures during the input-to-output conversion. To test this idea, we built a network model of feedforward inhibition (**Fig. 7A**)^23^. To simulate the modulation of CCK^+^-INs by noradrenaline, we depolarized the resting membrane potential of the interneurons in the model by 10 mV to match our electrophysiological results. This modulation facilitated the activation of the interneurons by excitatory inputs from presynaptic axons and enhanced the disynaptic feedforward inhibition generated by these interneurons (**Fig. 7B**–**D**, P < 0.05, n = 6 models).

**Figure 7.**
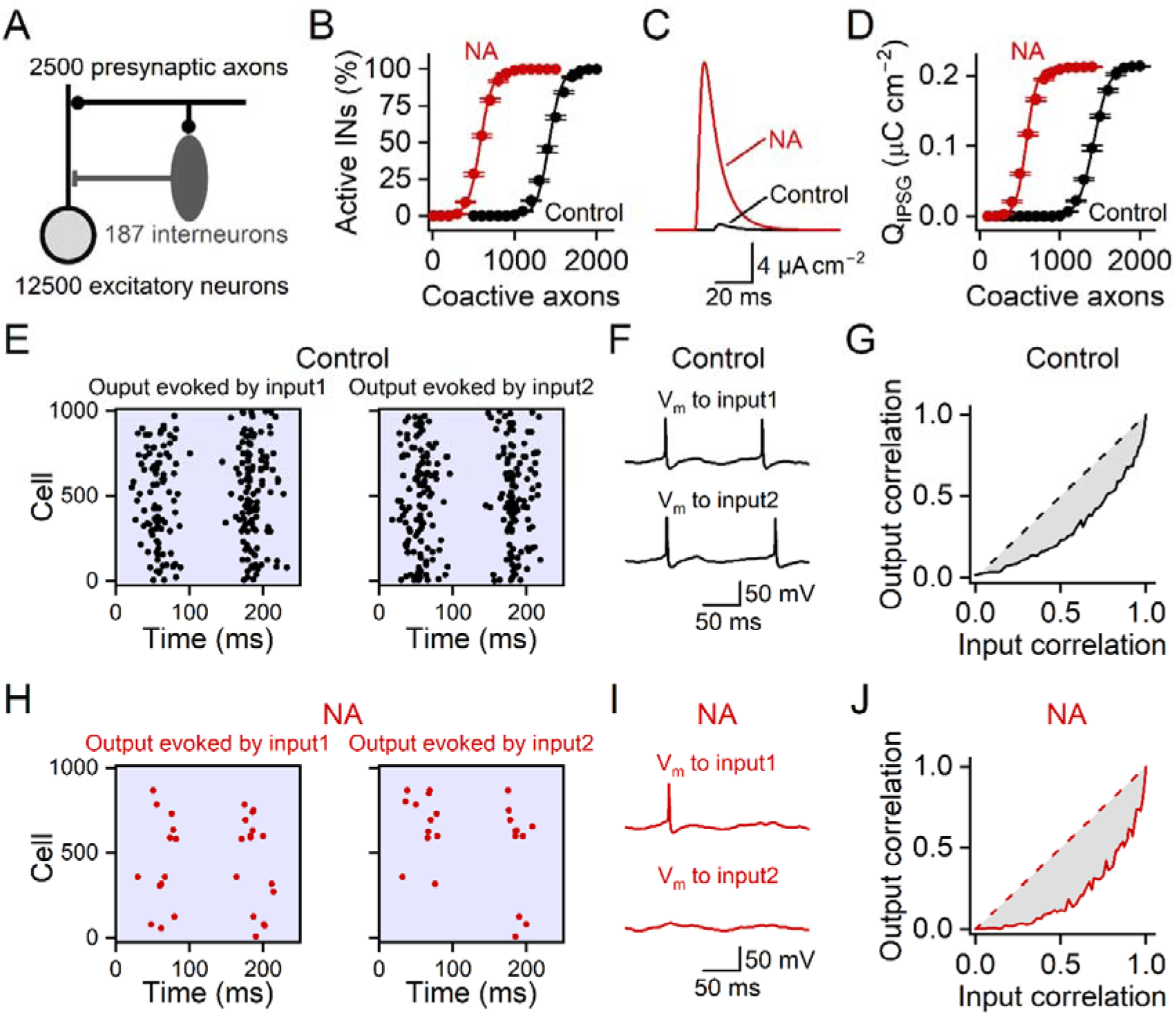
Facilitation of feedforward inhibition enhances decorrelation. (**A**) The structure of the network model, showing 2500 presynaptic axons that project to 12500 excitatory and 187 inhibitory neurons in the postsynaptic network. (**B**) Input-output relationship of interneurons, showing that the noradrenergic modulation facilitated their synaptic activation. Each data point in the input-output relationship was determined by co-activating a defined number of presynaptic axons, with each active axon firing a single AP. Lines represent Boltzmann functions fit to the data points. (**C**) Mean of the disynaptic feedforward IPSCs in response to the co-activation of 1100 presynaptic axons. (**D**) Similar to **B** but plots the charge density of the disynaptic IPSC against the total number of co-active axons. Circles and error bars in **B** and **D** represent mean ± SEM of the values obtained from six different models. (**E**) Raster plots showing the responses of postsynaptic excitatory neurons to two different inputs (input1 and input2, correlation = 0.89) under control conditions. Each symbol represents an AP generated by a postsynaptic excitatory neuron. For illustrative purposes, only the activity of 1000 randomly selected neurons was plotted. (**F**) Membrane potentials of a postsynaptic excitatory neuron evoked by the two inputs. (**G**) The correlation between output signals plotted against the correlation between input signals (continuous line). Decorrelation performance was quantified by the deviation (grey) of the relationship from the unity line (dashed line)^47^. (**H**–**J**) Similar to **E**–**G**, but the resting membrane potential of the interneurons was depolarized to mimic the modulation by noradrenaline. Data in **E** and **H** are from the same population of cells, and data in **F** and **I** are from the same cell. Results depicted in **E**–**J** have been reproduced in 6 different models, indicating that noradrenaline improved the decorrelation performance by 30.8 ± 3.0 % (n = 6 models, P = 0.03, two-sided Wilcoxon signed-rank test).

To analyze how the network decorrelated neuronal activity, we compared the correlation between input signals against that between corresponding output signals^46, 47^. In our simulations, input signals were represented by APs in presynaptic axons, whereas output signals were represented by APs in postsynaptic excitatory neurons. For each comparison, we activated two different ensembles of axons to create a pair of input signals and recorded the corresponding output signals (**Supplementary Fig. 4**). We found that the noradrenergic modulation made the output signals sparser and their patterns less correlated when compared to those under control conditions (**Fig. 7E**, **F**, **H**, **I**).

To quantify how well the network could perform decorrelation, we systematically changed the correlation between input signals and plotted it against the correlation of corresponding output signals. We determined the decorrelation performance based on the deviation of this relationship from the unitary line (**Fig. 7G**, **J**)^46, 47^. During the noradrenergic modulation, the deviation became more pronounced (**Fig. 7G**, **J**), indicating that facilitation of feedforward inhibition by noradrenaline could enhance the performance of the network to transform inputs into decorrelated AP outputs. These simulations led to the prediction that noradrenaline could facilitate the ability of granule cells to transform similar synaptic inputs into distinct AP outputs.

### Noradrenaline decorrelates granule cell activity patterns

To support the prediction with experimental evidence, we used electrophysiological and optical methods to compare granule cell AP patterns elicited by a pair of input signals with similar but distinct temporal structures^48^. Each input signal was generated by combining trains of excitatory synaptic inputs from two independent groups of medial perforant path axons, with the co-activation of these two groups of axons occurring at random time points (**Fig. 8A**–**B**). Given that these two input signals differed from each other in the timing of the co-activation of the two axonal pathways, and that granule cells are preferentially activated by highly synchronous excitatory inputs in noradrenaline, we predicted that noradrenaline could reduce the correlation between AP outputs evoked by these two input signals.

**Figure 8.**
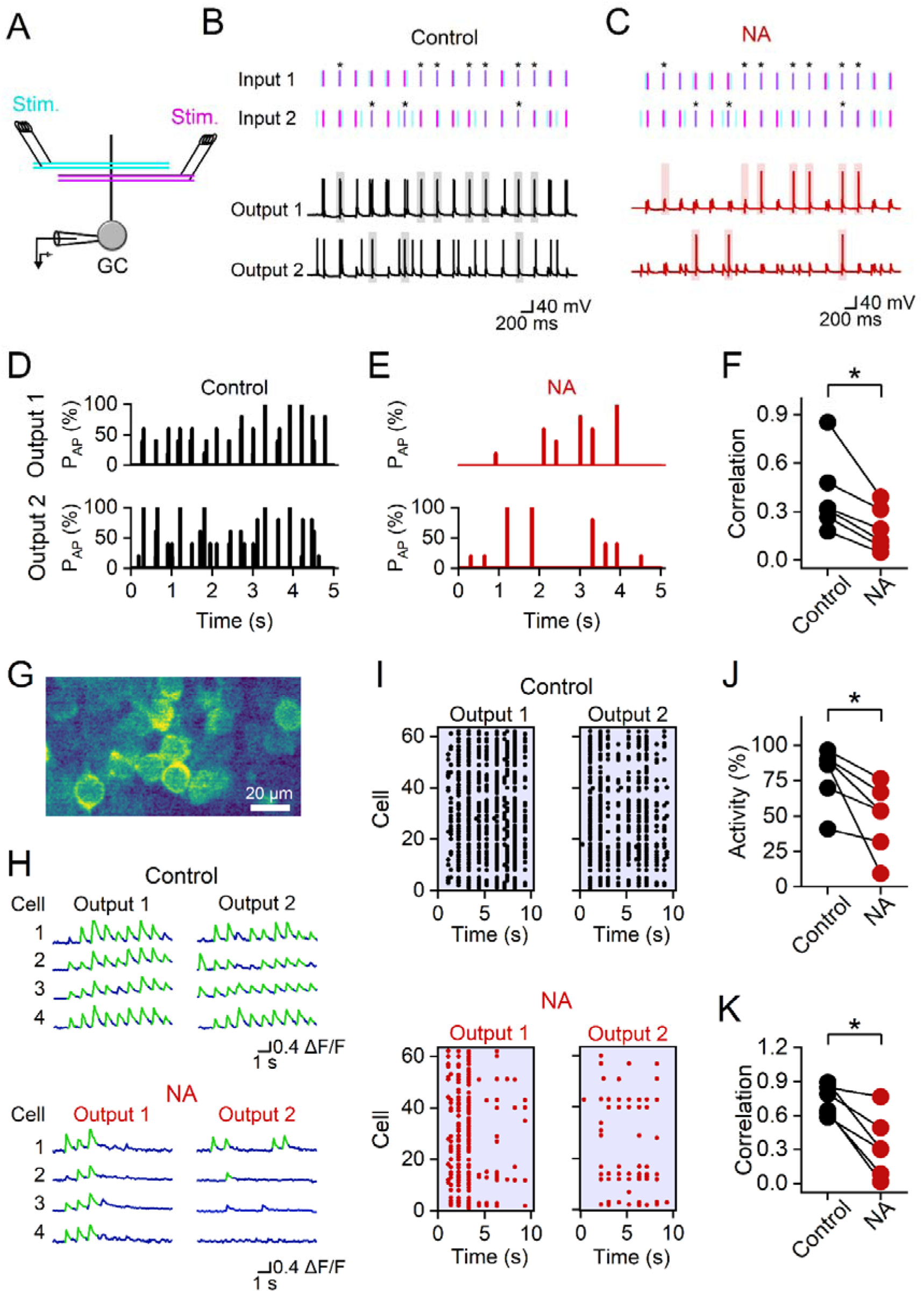
Noradrenaline decorrelates granule cell activity patterns. (**A**) Experimental design showing the stimulation of two independent groups of medial perforant path axons (in cyan and magenta) and electrophysiological recording of the responses in a granule cell. (**B**–**C**) Top, temporal pattern of the two input signals. Each of these input signals was generated by combining synaptic inputs from the two axonal pathways in **A**. Every dash symbol depicts the timing of a stimulus delivered to an axonal pathway (same color code as in **A**), which was modulated by a Poisson function (mean frequency = 3 Hz). Asterisks indicate random co-activation of both axonal pathways. Bottom, membrane potentials in a granule cell evoked by the two input signals under control conditions (**B**) and in bath-applied noradrenaline (**C**). Shaded areas highlight postsynaptic responses evoked by the co-activation of both axonal pathways. (**D**–**E**) The probability for the granule cell to generate APs in response to the pair of input signals, from the same experiment in **B**. The two input signals elicited similar postsynaptic responses under control conditions (**D**) but not in noradrenaline (**E**). (**F**) Summary graph showing correlation between granule cell activity patterns evoked by two input signals. (**G**) Confocal image of GCaMP8s-expressing granule cells. (**H**) Ca^2+^ signals in 4 randomly selected granule cells evoked by two input signals under control conditions (top) and in noradrenaline (bottom). Suprathreshold Ca^2+^ events are highlighted in green. (**I**) From the same experiment in **G**–**H**, showing suprathreshold Ca^2+^ events in 62 granule cells. Cells that did not generate any suprathreshold Ca^2+^ signals under control conditions were rejected from the analysis. (**J**) Noradrenaline inhibited the activity of the granule cell population. The activity was calculated by averaging the probability of synaptic stimuli to activate a granule cell across the granule cell population in the train. (**K**) Correlation of granule cell activity patterns at the population level, evoked by the two input signals. * indicates that P = 0.03 from 6 separate experiments. In summary plots, data points from the same experiment are connected by a line.

Indeed, patch-clamp recording from individual granule cells confirmed that AP patterns evoked by the two input signals became less correlated in noradrenaline when compared to those under control conditions (Output 1 and Output 2 in **Fig. 8B**– **F**). Detailed analyses of EPSP−AP coupling revealed that APs evoked by the co-activation of both axonal pathways were largely preserved in noradrenaline (probability to evoke an AP in the granule cell = 97.3 ± 1.0% under control conditions and 75.4 ± 4.4 % in noradrenaline; P = 0.03, two-sided Wilcoxon-signed rank test, n = 6 recordings). In sharp contrast, spikes evoked by the activation of a single axonal pathway were strongly inhibited following the application of noradrenaline (probability to evoke an AP in the granule cell = 68.4 ± 3.8 % under control conditions and 17.0 ± 4.7 % in noradrenaline, respectively; P = 0.03, two-sided Wilcoxon signed-rank test, n = 6 recordings). Thus, the decorrelation of granule cell output signals by noradrenaline is accompanied by a sparsening of their activity^13^.

To determine whether this result could be generalized to the granule cell population, we optically recorded Ca^2+^ signals in granule cells after virally expressing GCaMP8s in the dentate gyrus (**Fig. 8G**). Consistent with the electrophysiological recordings from single cells, suprathreshold Ca^2+^ signals in granule cells, which were evoked by the two input signals, became less frequent and less correlated following the application of noradrenaline (**Fig. 8H–K**). Collectively, these electrophysiological and optical data provide empirical evidence to confirm that noradrenaline can make granule cell APs sparser and their patterns decorrelated.

## Discussion

The dentate gyrus is densely innervated by noradrenergic axons originating from the locus coeruleus^17^. Behavioral studies have found that activation of the noradrenergic system modulates memory encoding in the dentate gyrus^49, 50^. To determine the underlying mechanism, previous studies have analyzed the effect of noradrenaline on long-term synaptic plasticity in the dentate gyrus^51, 52^. However, whether the release of noradrenaline can rapidly reorganize neuronal network dynamics in the dentate gyrus, on a timescale comparable to attention, remains unclear.

Here, we show that noradrenaline reversibly suppresses granule cell activity via an enhancement of synaptic inhibition mediated by CCK^+^-INs, and that such an action decorrelates action potential patterns generated by granule cells. By analyzing synaptic interactions between different cell types, our experiments differ conceptually from previous studies that focused on the intrinsic excitability of a single type of neuron in isolation^51, 53, 54, 55^. With this approach, our experiments have identified CCK^+^-INs as a key organizer of dentate gyrus network dynamics following the release of noradrenaline.

CCK^+^-INs make important contributions to the total interneuron population in the dentate gyrus^25, 26^. However, due to the technical limitation that reduces the throughput of the functional analyses of CCK^+^-INs^33, 35^, whether this abundance of dentate CCK^+^-INs is matched by their functional importance has remained an interesting but open question. Our experiments provide the first evidence to indicate that feedforward inhibition mediated by CCK^+^-INs is a critical determinant of the sparseness of granule cell activity, particularly following the release of noradrenaline. This result is relevant for identifying the mechanism underlying a reduction of granule cell activity level during behavioral states with heightened attention and arousal^10, 11,12^. We further demonstrated that noradrenaline, by facilitating this feedforward inhibition, enhances the ability of granule cells to discriminate inputs based on their temporal structure. This result provides an explanation for why chemogenetic inhibition of CCK^+^-INs in the dentate gyrus impairs context discrimination in a contextual fear memory discrimination task^56^.

Our results change the prevailing view that fast feedforward inhibition in the dentate gyrus is dominated by the contribution from PV^+^-INs^27, 43^. PV^+^-INs in the dentate gyrus express a sequence of subcellular specializations, which convert excitatory inputs targeting PV^+^-IN dendrites to powerful inhibitory outputs from their axons with a very short delay^45^. This feature endows PV^+^-INs with the ability to generate fast feedforward inhibition to regulate both the timing and probability of granule cell APs in response to afferent inputs. In contrast, CCK^+^-INs may appear poorly suited for such an operation because their speed of input-to-output conversion is relatively slow^57^. Despite this, our experiments provide converging evidence to demonstrate the existence and functional consequence of CCK^+^-IN-mediated feedforward inhibition in the dentate gyrus (**Fig. 4**). This raises the question about the mechanism underlying the earlier onset of CCK^+^-IN-mediated feedforward inhibition relative to granule cell APs, which is required for the effective suppression of granule cell firing by feedforward inhibition. One likely explanation lies in the difference in intrinsic excitability between granule cells and dentate CCK^+^-INs. The resting membrane potential of granule cells is substantially more negative than that of CCK^+^-INs (–79.6 ± 1.8 mV in granule cells, n = 7 cells; –62.0 ± 0.9 mV for CCK^+^-INs, n = 13 cells, P < 0.001), although the voltage threshold of APs between these two types of neurons does not differ (–40.6 ± 1.9 mV in granule cells, n = 6 cells; –43.8 ± 1.7 mV in CCK^+^-INs, n = 13 cells, P = 0.37). This difference in the resting membrane potential is enlarged by noradrenaline, which depolarizes CCK^+^-INs. Furthermore, granule cells display a larger membrane time constant than CCK^+^-INs (27.8 ± 3.0 ms in granule cells, n = 7 cells; 19.5 ± 1.2 ms for CCK^+^-INs, n = 13 cells, P = 0.02), although their input resistance is not significantly different from that of CCK^+^-INs (242.3 ± 38.2 MΩ for granule cells, n = 7 cells; and 215.8 ± 16.4 MΩ for CCK^+^-INs, n = 13 cells, P = 0.8). The combination of these factors delays AP initiation in granule cells in response to afferent inputs, thereby creating a broad temporal window for the recruitment of CCK^+^-IN-mediated feedforward inhibition.

Our experiments have identified CCK^+^-INs as important intermediaries between the release of noradrenaline and the subsequent inhibition of granule cell firing, and they raise an interesting question about the mechanism underlying the noradrenergic upregulation of CCK^+^-IN excitability. CCK^+^-INs in the dentate gyrus express adrenergic receptors^58^. Furthermore, electrophysiological experiments have found that noradrenaline can depolarize CCK^+^-INs in the frontal cortex in the presence of glutamate receptor blockers^59^. Collectively, these two studies suggest that noradrenaline may directly modulate the intrinsic excitability of CCK^+^-INs by activating adrenergic receptors on their membrane. In addition, NA may modulate astrocytic signaling in the dentate gyrus^58, 60^. Because astrocytes in the dentate gyrus express adrenergic receptors, noradrenaline, in essence, can indirectly modulate neuronal excitability by triggering the release of gliotransmitters from astrocytes^61, 62^. Future experiments, which specifically inhibit the function of astrocytes, are needed to test this idea.

How does the release of noradrenaline in the dentate gyrus contribute to memory formation in the hippocampus? Dentate gyrus granule cells represent a major source of excitatory inputs to the CA3 network during memory formation^2^. When compared to the CA1 and the dentate gyrus, a distinct feature of the CA3 neuronal network is the recurrent excitatory synaptic connections between pyramidal neurons in this hippocampal subregion^63^. While this anatomical feature is critical for the rapid association of different components of an episodic memory during learning^3, 64, 65^, it also renders memory representations susceptible to the interference brought about by correlated neuronal activity^6, 9^. Consequently, decorrelation of granule cell activity has strong implications for memory functions performed by the downstream CA3 network^1, 2, 3, 6^. Our results, together with previous studies, suggest that release of noradrenaline in the dentate gyrus may contribute to this operation in two complementary ways. First, the enhancement of feedforward inhibition by noradrenaline allows granule cells to be preferentially activated by highly synchronous excitatory synaptic inputs. This acute effect, which occurs rapidly following the release of noradrenaline, provides individual granule cells with the ability to generate decorrelated AP patterns by identifying subtle differences in the temporal structure of excitatory inputs from the entorhinal cortex^66^. Second, the facilitation of long-term potentiation in granule cells by noradrenaline^51, 52^, when paired with lateral inhibition, implements a competitive learning mechanism that makes granule cell activity more specific at the single-cell level and decorrelated at the population level during longitudinal learning over consecutive days^67, 68^. Collectively, these two mechanisms, which operate on different temporal scales, may facilitate memory storage in the CA3 and subsequent retrieval.

In addition, our experiments reveal that noradrenaline promotes the decorrelation of granule cell activity at both single-cell and population levels, revealing a novel mechanism by which noradrenaline contributes to high-order cognitive functions beyond its established role in neuronal gain and plasticity^51, 69^. Correlations within neuronal population activity have a strong influence on information encoding, rendering the mechanisms governing correlated and decorrelated firing patterns critical for neuronal coding^6^. Several mechanisms, including expansion recoding based on Marr’s theory in the cerebellum and neuronal assembly competition using recurrent feedback synaptic connections, have been proposed to account for the decorrelation performed by the dentate gyrus network^1, 6, 46, 47, 70^. In contrast, our results demonstrate that noradrenergic enhancement of feedforward inhibition implements a distinct mechanism to facilitate decorrelation in the dentate gyrus. Notably, the shorter temporal delay of disynaptic feedforward inhibition, when compared to decorrelations using polysynaptic recurrent synaptic connections, is expected to improve the discrimination of inputs with a finer temporal structure, as well as increase the throughput of the decorrelation operation. Given the widespread expression of CCK^+^-INs, and the abundance of noradrenergic inputs and feedforward inhibitory motifs in the cortex, we speculate that our findings can be generalized to other cortical areas.

## Materials & Methods

### Animals

Experiments were ethically approved by the Norwegian Food Safety Authority (Mattilsynet) and performed in strict accordance with institutional, national, and European guidelines for animal experimentation.

All transgenic mouse lines were obtained from The Jackson Laboratory, and the following mouse lines were used in this study: hemizygous TH-cre mice (strain: 008601), heterozygous PV-IRES-cre mice (strain: 017320), heterozygous CCK-IRES-cre mice (strain: 012706), heterozygous GAD2-IRES-cre (stain: 010802) and Ai14 (strain: 007914) mice, and wild-type C57BL/6 mice. Hemizygous TH-cre mice were generated by pairing hemizygous parents, heterozygous GAD2-IRES-cre and Ai14 mice were generated by pairing homozygous GAD2-IRES-cre mice with Ai14 mice. Other lines were generated by crossbreeding homozygous mice with wild-type mice. In addition, we labeled PV^+^-INs by crossbreeding homozygous PV-IRES-cre mice with homozygous Ai14 reporter mice.

Pharmacological experiments were performed in acute slices prepared from the brains of male and female juvenile mice (3-to 8-week-old), whereas optogenetic, chemogenetic and GCaMP imaging experiments were performed in acute brain slices prepared from adult mice (3-to 9-month-old). To ensure that our results were not confounded by age, we performed additional cell-attached recordings in granule cells from adult wild-type mice. A correlation analysis confirmed that the inhibition of granule cell activity by bath-applied noradrenaline was age-independent (Spearman ρ = 0.23, P = 0.37, n = 17 mice).

### Viral injections

All viruses were purchased from Addgene (**Supplementary Table 1**). For optogenetic stimulation and labeling of noradrenergic axons, we used pAAV-Syn-FLEX-rc[ChrimsonR-tdTomato]. For the optical recording of granule cell activities, we used pGP-AAV-syn-jGCaMP8s-WPRE and pAAV-mDlx-NLS-mRuby2. For chemogenetic inhibition of PV^+^-INs, we injected pAAV-hSyn-DIO-hM4D(Gi)-mCherry into the hippocampus of PV-IRES-cre mice. For labeling CCK^+^-INs, we used pAAV-hDlx-Flex-GFP.

Stereotaxic viral injections were performed in adult mice (3-to 8-month-old). During surgery, animals were kept under isoflurane anesthesia (3% for induction, 1– 2% for maintenance) and local analgesia (subcutaneous injection of 0.25% Marcaine), and body temperature was maintained at 37°C. Viruses were delivered to the target areas using a Nanoject III injector (Drummond Scientific) at a speed of 1 nL per second. Titer and stereotaxic coordinates are provided in **Supplementary Tables 1** and **2**. Postoperative analgesia (Temgesic, 0.1 mg per kg) was administered subcutaneously for up to 3 days.

Optogenetic experiments were performed 4–5 weeks following viral injections, whereas other experiments were performed 2–3 weeks post-surgery. The success of viral gene expression was functionally confirmed by recordings of light-evoked APs in noradrenergic axons (**Fig. 1D**–**F**) and GCaMP8s signals in granule cells (**Figs. 2F**–**I**, **8G**–**K** and **Supplementary Fig. 1**). The effectiveness of chemogenetic inhibition of PV^+^-INs was confirmed by cell-attached recordings from these interneurons, which indicated robust suppression of PV^+^-IN activity by bath-applied CNO (PV^+^-IN AP probability in response to medial perforant path inputs: 79 ± 8.5% under control conditions and 19 ± 10.9% in 1 µM CNO, respectively. P = 0.03, n = 6 experiments in slices from 5 mice, two-sided Wilcoxon signed-rank test).

To target CCK^+^-INs for patch-clamp recordings, we performed retro-orbital injection of pAAV-hDlx-Flex-GFP into CCK-IRES-Cre mice at postnatal day 1 under isoflurane anesthesia. After the surgery, animals were returned to a heating pad for recovery before being transferred back to the home cage. Electrophysiological recordings from individual GFP-labeled interneurons were performed 3–5 weeks after the retro-orbital injection.

### Electrophysiological recordings

Slices prepared from juvenile mice were cut in an ice-cold, sucrose-containing extracellular solution using a vibratome (VT1200, Leica Microsystems), incubated in a storage chamber filled with the standard physiological extracellular solution at ∼34°C for 30 minutes, and subsequently stored at room temperature. In contrast, slices prepared from adult mice were cut in an ice-cold, HEPES-based extracellular solution and subsequently incubated in a choline-rich incubation solution at ∼34°C for 10 minutes before being transferred to a storage chamber filled with the standard physiological extracellular solution^71^.

Prior to recordings, slices were individually transferred into a recording chamber and superfused with the standard physiological extracellular solution. Electrophysiological recordings were performed at near-physiological temperature (∼32°C; range: 31–33°C). In each experiment, the temperature was maintained at a constant level within ± 0.5°C. Recordings were obtained using an Olympus BX51 infrared differential interference contrast (IR-DIC) microscope. Fluorescently labeled interneurons were identified with an Andor DSD2 confocal unit (Oxford Instruments).

Pipettes for patch-clamp recordings were fabricated from thick-walled borosilicate glass capillaries (outer diameter: 2 mm, inner diameter: 1.16 mm) with a horizontal pipette puller (P-97, Sutter Instruments). When filled with the internal solution, they had a resistance of 5–11 MΩ for cell-attached recordings and 5–9 MΩ for whole-cell recordings.

Cell-attached recordings (seal resistance > 3 GΩ) were performed at the soma of identified neurons by voltage-clamping the electrode at –70 mV. Pipettes for these recordings were either filled with the standard physiological external solution or the standard K-gluconate-based internal solution. In a subset of cell-attached recordings, 50 µM Alexa Fluor 488 was included in the pipette solution, and the recorded neuron was examined with epifluorescence illumination at the end of the experiment. None of the cells displayed cytoplasmic Alexa Fluor 488 signals, indicating that the integrity of the cell-attached configuration was maintained throughout the experiment.

To perform axonal loose-patch recordings^72^, we used the Andor DSD2 confocal unit to search for tdTomato-labeled noradrenergic axons in the dentate gyrus. After correlating the confocal image with the IR-DIC image from the same field of view, an axonal recording was obtained. Axonal patch electrodes had a resistance of ∼15 MΩ when filled with the standard internal solution. Throughout the recording, small negative pressure (< 20 mbar) was applied to the inside of the pipette to prevent the leakage of the pipette solution into the bath and to maintain the seal resistance (0.2– 1 GΩ). Throughout the experiment, the recording pipette was voltage-clamped at 0 mV.

Current-and voltage-clamp whole-cell recordings were performed using a Multiclamp 700B amplifier (Molecular Devices). Series resistance in current-clamp recordings was 7–19 MΩ. Pipette capacitance compensation in current-clamp recordings was performed by adjusting the pipette capacitance neutralization function of the amplifier, and bridge balance was monitored and readjusted as required during each experiment. Voltage signals in current-clamp recordings were obtained at the resting membrane potential, except for the comparison of input resistance under control conditions with that in noradrenaline. In these experiments, input resistance in noradrenaline was measured after the somatic membrane potential was restored to the value under control conditions by somatic current injection. In voltage-clamp recordings, the series resistance was 9.0–18.3 MΩ, and series resistance compensation was not performed. Individual voltage-clamp experiments were discarded if the series resistance in the experiment changed by more than 20%.

Recordings of local field potentials were performed with a Multiclamp 700B amplifier, using a glass recording electrode (pipette resistance: 1–3 MΩ when filled with the standard physiological extracellular solution) positioned in the granule cell layer.

Signals were low-pass filtered at 10 kHz in current-clamp recordings, at 4–10 kHz in voltage-clamp recordings and extracellular (cell-attached, loose patch, and field) recordings. The filtered signals were sampled at 50 or 100 kHz with a Digidata 1322 converter board (Molecular Devices). Pulse protocols were generated using pClamp 9 or 10 (Molecular Devices).

To stimulate medial perforant path axons, we positioned a bipolar tungsten stimulation electrode in the middle third of the molecular layer. To avoid contamination by monosynaptic IPSCs, the extracellular stimulation electrode and the recording electrode were positioned in separate blades of the dentate gyrus, i.e. the supra- and infrapyramidal blades, except for recordings of monosynaptic IPSCs. Synaptic stimulation (duration: 100 µs) was delivered once every 10 s. In cell-attached recordings from granule cells, we adjusted the intensity of the synaptic stimulation to evoke an AP with a probability of 67.6 ± 2.4% under control conditions (n = 71 experiments). In recordings of population spikes and monosynaptic currents, the stimulation intensity was adjusted to elicit responses with an amplitude that was approximately half of the maximum. In whole-cell recordings of disynaptic IPSCs, cell-attached recordings from interneurons, and optical records of Ca^2+^ signals in granule cells, we simultaneously measured the activity of nearby granule cells with a separate electrode, and the synaptic stimulation intensity was calibrated based on the granule cell activity using the criterion described above. When we determined the effect of noradrenaline on the granule cell response to repetitive synaptic stimulation (**Supplementary** Fig. 3), the synaptic stimulation intensity was adjusted so that high-frequency bursts (> 150 Hz) represent approximately 5% of all postsynaptic APs evoked by the synaptic stimulation protocol^12^. Synaptic currents were recorded at – 50 mV. Monosynaptic IPSCs in granule cells were isolated with bath-applied DNQX (10 µM) and DL-APV (50 µM). EPSCs were isolated with bath-applied SR95531 (4 µM).

To determine the effect of noradrenaline on uIPSCs, we performed paired recordings from monosynaptically connected presynaptic interneurons and postsynaptic granule cells. Series resistance in the postsynaptic neuron was 7–11 MΩ and remained stable in each experiment. Presynaptic APs were evoked from the interneuron resting membrane potential, and the postsynaptic granule cell was voltage-clamped at –70 mV without series resistance compensation.

### Optogenetic stimulation of noradrenergic axons

To induce noradrenaline release in acute hippocampal slices with an optogenetic approach, light pulses (duration: 10 ms, wavelength: 590 nm) were generated with a collimated LED light source (Thorlabs) for wide-field illumination. The output power was 5.2–8.6 mW mm^−2^ at the front focal plane of the microscope objective unless otherwise specified. Loose-patch recordings confirmed that a 10-ms light pulse at this output power was sufficient for activating ChrimsonR-expressing noradrenergic axons in the dentate gyrus (**Fig. 1E**, n = 8 axons).

The firing pattern of noradrenergic neurons changes in a behavioral state-dependent manner^73^. With the cell-attached method, we found that stimulating noradrenergic axons with a train of five light pulses at 10 Hz (delivered 100 ms before the electrical stimulation of medial perforant path axons and repeated every 10 s for 5 minutes) reliably inhibited granule cell activity (**Fig. 1H**–**1J**). In sharp contrast, reducing the frequency of the light pulses in the train to 3 Hz was insufficient for inhibiting granule cells (probability of activating granule cells by inputs from medial perforant path axons = 68.0 ± 8.0% under control conditions and 80.0 ± 15.5% following optogenetic activation of noradrenergic axons, P = 0.46, n = 5 cells, two-sided Wilcoxon signed-rank test). Consequently, we used the 10-Hz photo-stimulation protocol for the optogenetic experiments illustrated in **Figs**. **1**, **2**, and **5**.

### Identification of interneurons

The identity of recorded interneurons was confirmed based on their electrophysiological and morphological properties. The electrophysiological properties of interneurons were characterized with a whole-cell pipette that contained 0.2% biocytin. When we performed cell-attached recordings from interneurons, they were routinely repatched with a whole-cell pipette containing biocytin at the end of the experiment for the characterization of their AP phenotype and subsequent histological analyses. Following electrophysiological recordings, slices were fixed overnight in 4% paraformaldehyde and subsequently incubated in PBS containing 0.3% Triton X-100 for 2 hours. Morphology of biocytin-filled cells was revealed with Alexa 594-conjugated streptavidin (1:1000, Thermo Fisher Scientific) and captured with an Andor Dragonfly confocal microscope (Oxford Instruments).

To fluorescently label PV^+^-INs for patch-clamp recordings, we crossbred homozygous PV-IRES-Cre mice with homozygous Ai14 reporter mice. We prepared acute hippocampal slices from the heterozygous offspring and obtained recordings from tdTomato-expressing interneurons in the dentate gyrus. The identity of these neurons was further confirmed by the non-accommodating, fast-spiking phenotype (steady-state AP frequency > 150 Hz at near-physiological temperature in response to 1-s, 0.3- to 1-nA somatic current pulses), and by the morphological properties of the axonal arbor, which was largely restricted to the granule cell layer (**Supplementary Fig. 2A–C**).

To obtain recordings from CCK^+^-INs, we performed retro-orbital injection of AAV9-hDlx-Flex-GFP in CCK-IRES-Cre mice^35, 36^. We targeted GFP-expressing neurons for electrophysiological recordings in acute hippocampal slices prepared from these mice 3–5 weeks after the injection. Because this intersectional approach may create unspecific GFP expression in PV^+^-INs and neurogliaform cells^33^, we used a combination of functional and morphological criteria to avoid sampling from these two interneuron subpopulations. We excluded PV^+^-INs based on their stereotypical fast-spiking phenotype and dense axonal projection to the granule cell layer. Neurogliaform cells were identified by delayed AP initiation in response to the injection of a 1-s depolarizing current pulse, relatively short dendrites, and a dense axon plexus surrounding their soma^74^. After excluding putative PV^+^-INs and neurogliaform cells, our CCK^+^-IN sample included 16 hilar commissural-associational pathway-associated (HICAP) cells, 5 total molecular layer (TML) cells, and 4 cells with incomplete axonal morphology that prevented further classification. Dendrites of these interneurons were aspiny, distinguishing our CCK^+^-IN sample from granule cells and mossy cells. A hallmark of CCK^+^-INs is the expression of CB1 receptors on their axonal terminals^75^. To confirm the specificity of our CCK^+^-IN samples, we determined the effect of ACEA (500 nM) on the synaptic output generated by cells randomly selected from our samples. In five monosynaptically connected presynaptic CCK^+^-IN and postsynaptic granule cell pairs, ACEA consistently blocked uIPSCs generated by the interneuron (uIPSC charge = 1.70 ± 0.58 pC and 0.08 ± 0.03 pC under control conditions and in ACEA, respectively, P = 0.04, n = 5, two-sided Wilcoxon signed-rank test).

### Optical recordings of granule cell activity

Optical recordings of Ca^2+^ signals in **Fig. 2** and **Supplementary** Fig. 1 were performed in acute hippocampal slices prepared from wild-type mice, 2–3 weeks after injecting pGP-AAV-syn-jGCaMP8s-WPRE and pAAV-mDlx-NLS-mRuby2 into the dentate gyrus of wild-type mice. Because the expression of mRuby2 was driven by an interneuron-specific enhancer^36^, mRuby2-expressing neurons were classified as GABAergic interneurons and excluded from subsequent analyses. Experiments in **Fig. 8G–K** were performed after injecting pGP-AAV-syn-jGCaMP8s-WPRE into the dentate gyrus of heterozygous GAD2-IRES-cre and Ai14 mice. Interneurons were identified based on the expression of tdTomato and excluded from subsequent analyses. To exclude mossy cells, analyses were restricted to neurons with a small soma in the granule cell layer.

Optical recordings were performed with an Olympus BX51 microscope equipped with the Andor DSD2 confocal unit at near-physiological temperature (∼32°C). Ca^2+^ signals in granule cells were evoked by stimulating medial perforant path axons with a bipolar stimulation electrode. GCaMP8s was excited at a wavelength of 470 nm with an LED light source, and the emitted fluorescence was collected with an Olympus water immersion objective (NA 0.9). GCaMP8s signals were isolated with a 525/45 nm emission filter and recorded at a sample rate of 20 Hz using a Zyla 4.2 sCMOS camera (Oxford Instruments).

Optical signals were processed and analyzed with ImageJ (NIH). We manually selected regions of interest (ROIs) from granule cell somata, and background fluorescence was subtracted from fluorescent signals extracted from the ROIs. To calculate the Ca^2+^-dependent increase in fluorescence (ΔF/F_0_) in each experiment, we computed F_0_ based on fluorescent signals in a 5-s time window at the beginning of each recording. Suprathreshold Ca^2+^ signals were identified as those with a peak ΔF/F_0_ exceeding 0.24. We established this criterion by correlating optically recorded Ca^2+^ signals with electrophysiologically identified APs in a separate set of experiments (**Supplementary** Fig. 1). To quantify the fraction of granule cells activated by the synaptic stimulation (**Fig. 2I**), we divided the number of granule cells activated in response to the stimulation by the total number of granule cells. Granule cells that failed to produce any suprathreshold response throughout the control period were excluded from the analyses of **Figs. 2** and **8**.

### Analysis of decorrelation of granule cell AP patterns

To determine how granule cells transform different synaptic input signals into decorrelated AP patterns (**Fig. 8**), we delivered a pair of input signals to granule cells and compared the corresponding granule cell AP patterns. Each input signal was generated by repetitively stimulating two independent groups of medial perforant path axons. The timing of the synaptic stimuli was modulated by a Poisson function, causing the co-activation of both axonal pathways to occur at random time points. The mean frequency of the Poisson function was 3 Hz in experiments depicted in **Fig. 8B**–**F**, matching the AP frequency range of neurons in superficial layers of the medial entorhinal cortex during spatial navigation^66^. When comparing the two input signals, the timing of stimuli delivered to one group of medial perforant path axons was identical, whereas the timing of stimuli delivered to the other group was different. In experiments illustrated in **Fig. 8G**–**K**, the frequency of the Poisson function was reduced to 1 Hz to compensate for the low temporal resolution when using Ca^2+^ signals to assess neuronal activity. To ensure that the enhancement of decorrelation by noradrenaline did not depend on a specific set of input signals, a new pair of input signals was generated before each experiment. The pair of input signals was delivered to the slice five times under control conditions and five times in bath-applied noradrenaline in every experiment. After recording granule cell output signals evoked by these input signals, we constructed peristimulus time histograms (PSTHs) based on the probability of individual synaptic stimuli to evoke a suprathreshold response. The duration of the temporal bin in the PSTH is 5 ms in **Fig. 8D**–**F**. A control analysis using temporal bins with different durations (range: 1–100 ms) produced qualitatively similar results. In the analysis of **Fig. 8K**, the duration of the temporal bin was increased to 275 ms to compensate for the low temporal resolution of Ca^2+^ signals. Granule cell APs in **Fig. 8B**–**F** were identified based on the recording of membrane potentials, whereas suprathreshold Ca^2+^ signals in **Fig. 8G**–**K** were identified based on the criterion in **Supplementary** Fig. 1. The correlation between output signals in each single granule cell was subsequently quantified by comparing these PSTHs, and the correlation of output signals generated by the granule cell population represents the mean correlation value from all granule cells included in the analysis. Decorrelation of postsynaptic responses was determined by comparing the correlation under control conditions with that in bath-applied noradrenaline.

### Solutions and chemicals

The standard physiological extracellular solution contained 125 mM NaCl, 25 mM NaHCO_3_, 2.5 mM KCl, 1.25 mM NaH_2_PO_4_, 2 mM CaCl_2_, 1 mM MgCl_2_, 25 mM D-glucose, and 0.1 mM ascorbic acid (equilibrated with a 95% O_2_ and 5% CO_2_ gas mixture). The sucrose-based saline for preparing slices from juvenile mice contained 87 mM NaCl, 25 mM NaHCO_3_, 25 mM glucose, 75 mM sucrose, 2.5 mM KCl, 1.25 mM NaH_2_PO_4_, 0.5 mM CaCl_2_, and 7 mM MgCl_2_. The HEPES-based extracellular solution for preparing slices from adult mice contained 92 mM NaCl, 2.5 mM KCl, 1.2 mM NaH_2_PO_4_, 20 mM HEPES, 30 mM NaHCO_3_, 25 mM glucose, 5 mM sodium ascorbate, 2 mM thiourea, 3 mM sodium pyruvate, 12 mM N-acetyl-L-cysteine, 10 mM MgSO_4_, and 0.5 mM CaCl_2_, pH adjusted to 7.3 with NaOH. The choline-rich incubation solution for slices from adult mice contained 92 mM choline chloride, 2.5 mM KCl, 1.2 mM NaH_2_PO_4_, 30 mM NaHCO_3_, 20 mM HEPES, 25 mM glucose, 5 mM sodium ascorbate, 2 mM thiourea, 3 mM sodium pyruvate, 12 mM N-acetyl-L-cysteine, 10 mM MgSO_4_, and 0.5 mM CaCl_2_.

The standard internal solution for whole-cell recordings contained 140 mM K-gluconate, 5 mM NaCl, 0.5 mM EGTA, 2 mM MgCl_2_, 2 mM Na_2_ATP, 0.4 mM Na_2_GTP, 10 mM HEPES, pH adjusted to 7.3 with KOH. In paired recordings from monosynaptically connected neurons, the presynaptic intracellular solution contained 135 mM K-gluconate, 20 mM KCl, 0.1 mM EGTA, 2 mM MgCl_2_, 2 mM Na_2_ATP, 0.4 mM Na_2_GTP, 0.2% biocytin, and 10 mM HEPES, whereas the postsynaptic intracellular solution contained 145 mM KCl, 0.1 mM EGTA, 2 mM MgCl_2_, 2 mM Na_2_ATP, 0.4 mM Na_2_GTP, 10 mM HEPES, and 2 mM QX 314.

### Data analysis

Analyses were performed using Clampfit 9 and 10 (Molecular Devices), Origin 2022 (Microcal), SPSS Statistics (IBM), and Matlab R2018a (MathWorks).

To determine the probability for medial perforant path inputs to evoke an AP in a recorded neuron, the synaptic stimulation protocol was repeated for a minimum of 5 times, and the probability was quantified based on the total number of evoked APs in these trials. In other illustrated time courses, each data point represents the mean value determined from a minimum of 5 consecutive trials. To quantify the size of a population spike^76^, we fit the two positive peaks flanking the population spike with a linear function and calculated the integral between the linear function and the population spike. To determine the AP phenotype of interneurons, we injected a series of 1-s depolarizing current pulses into the soma. The steady-state AP frequency was computed from the total number of APs during the 1-s train. To determine the input resistance of individual neurons, we injected a series of 1-s current pulses of various amplitudes into the soma. After plotting the amplitude of the steady-state voltage response against that of the corresponding current pulse in the subthreshold range, we fit the voltage-current relationship with a linear function. The input resistance was determined from the slope of the linear function.

To determine the reversal potential of the noradrenaline-sensitive synaptic current, we recorded this current at membrane potentials ranging from –50 mV to –90 mV. After plotting the peak amplitude of the current against the corresponding membrane potential, we fit the current-voltage relationship with a linear function. We estimated the reversal potential of the current by finding where the linear function intercepted the abscissa, which represented the membrane potential. The onset of synaptic currents was defined as the time point where the amplitude of these currents exceeded three times the standard deviation of the background noise, which was computed from a 70-ms time window immediately before synaptic stimulation. The timing of APs in cell-attached recordings was quantified by using the time point at the trough of the action current. For illustrative purposes, synaptic stimulation artifacts were manually truncated in the depicted cell-attached recordings and removed from all other depicted electrophysiological traces.

### Simulations of feedforward inhibition and decorrelation

The network model consisted of a presynaptic component and a postsynaptic component (**Fig. 7A**). The presynaptic component consisted of 2500 excitatory axons, whereas the postsynaptic component had 12500 excitatory principal neurons. This ratio of pre-to postsynaptic neurons matches that of projection neurons from layer 2 of the entorhinal cortex to dentate granule cells^6, 77^. In addition, the postsynaptic component had 187 regular-spiking inhibitory interneurons to capture the contribution of CCK^+^-INs (approximately 1.5%) to the dentate neuronal population^25, 78, 79^. Each postsynaptic neuron received excitatory inputs from 50 randomly selected presynaptic axons, corresponding to 2% of the total number of presynaptic axons^77^, and each interneuron inhibited 700 randomly selected excitatory neurons^80^. **Supplementary Table 3** provides a detailed description of the passive and active membrane properties of the postsynaptic neurons.

Each excitatory neuron in the postsynaptic network was represented by a multi-compartment ball-and-stick model, with a 210-µm-long apical dendrite connected to an isopotential soma. Excitatory synapses in excitatory neurons were randomly distributed to the middle third of their dendrites, whereas inhibitory synapses were assigned to a random location in the proximal two-thirds of the dendrite. The excitatory synapses contained both AMPA and NMDA conductances. A detailed description of synaptic conductances is provided in **Supplementary Table 4**.

Each interneuron in the postsynaptic network was represented by a single-compartment cell that received excitatory inputs from presynaptic axons. Excitatory synapses in these interneurons contained both AMPA and NMDA conductances (**Supplementary Table 4**). To analyze feedforward inhibition in isolation, interneurons did not receive recurrent feedback excitatory inputs from the postsynaptic excitatory neurons. Furthermore, neither inhibitory synapses nor gap junctions were implemented between interneurons in the model.

Excitatory synapses were activated by APs in presynaptic axons (mean steady-state AP frequency = 35.9 Hz). The AP pattern in each presynaptic axon was represented by all-or-none binary signals as a function of time, and the timing of individual presynaptic APs was determined by a random Poisson process. The output of the Poisson process was further modulated by an 8-Hz sinusoidal function to mimic theta modulation of excitatory synaptic inputs from the entorhinal cortex^41, 46^.

To analyze decorrelation, we created pairs of presynaptic activity patterns. For each pair, we first generated a reference signal (Input1) by randomly activating a defined number of axons. Subsequently, we created another signal (Input2) by replacing part of the active axon population with randomly selected axons that were inactive in the reference signal (**Supplementary** Fig. 4). This approach produced two input signals with various degrees of correlation. For each signal, we constructed a population vector to represent the steady-state AP frequency of every axon, which was determined from the number of APs in each axon in a 400-ms window^46^. We quantified the correlation between a pair of population vectors based on the cosine of the angle between them^13^. The same approach was used to calculate the correlation between output signals generated by the postsynaptic network. After plotting the correlation between output signals against that of corresponding input signals, we quantified the decorrelation performance of the network based on the integral between the unity line and the correlation relationship (**Fig. 7G** and **J**)^47^.

Simulations were performed using Neuron 8.0–8.2 running on a high-performance computing cluster. The simulation duration was 500–600 ms following the activation of presynaptic axons, and the integration time step in the simulations was 0.1 ms. For each combination of parameters, the simulation was performed in six different models under random initiation conditions.

### Quantification and statistical analysis

Membrane potentials are specified without correction for liquid junction potentials, except for the analysis of the reversal potential of noradrenaline-sensitive synaptic currents. Values and summary graphs indicate mean ± standard error of the mean (SEM) except for **Supplementary Fig. 1E.** Significance of the difference between two groups of data was assessed by a two-sided nonparametric Wilcoxon signed-rank test or Wilcoxon rank sum test. Differences were considered significant if P < 0.05. The difference between input-output relationships was assessed with a bootstrap method with a minimum of 10000 iterations (**Fig. 7B** and **D**).

## Supporting information

Supplementary Figures, Tables, and References

## Acknowledgments

We thank Drs. Ethan M. Goldberg, Fabian Roth, and Matthijs Dorst for critically reading a previous version of the manuscript. This project has received funding from the Norwegian Research Council, grant agreement numbers 250866, 276047, and 281252 (to H.H.).

## Data availability

Original data, code for the simulations, and analysis programs will be provided by the corresponding author (H.H.) upon request.

## Author contributions

I.G. and H.H. performed experiments and analyzed data. H.H. performed simulations and wrote the first draft of the paper. All authors jointly designed the experiments and revised the paper.

