## Supplementary Figures, Tables, and References for "Sparsening and Decorrelation of Granule Cell Activity in the Dentate Gyrus by Noradrenaline"

Iulia Glovaci et al.

Division of Physiology, Department of Molecular Medicine, Institute of Basic Medical Sciences, University of Oslo, Norway

**This PDF file includes:**

Supplementary Figures 1–4

Supplementary Tables 1–4

Supplementary reference

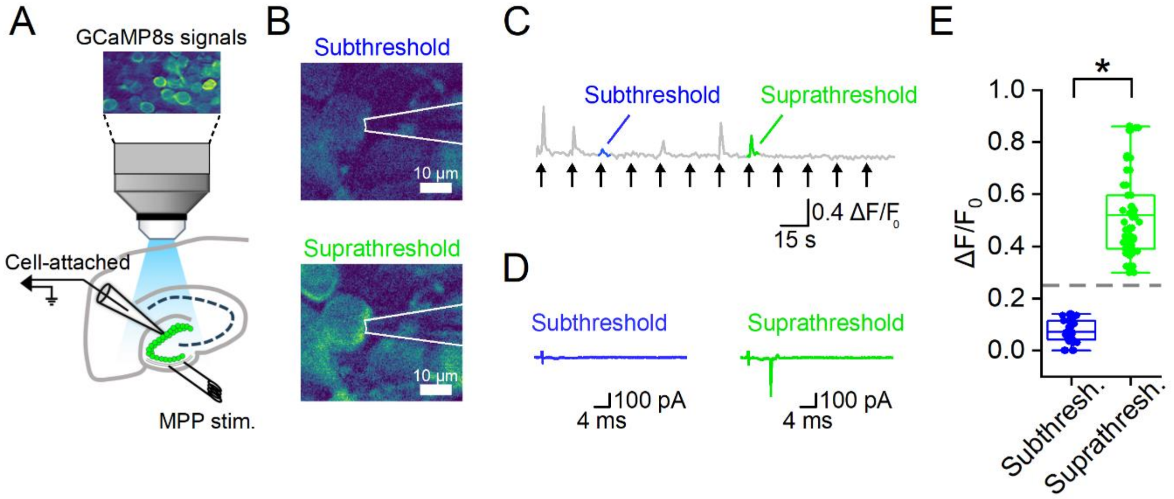


**Supplementary Figure 1. Separating sub- from suprathreshold $Ca^{2+}$ signals in individual granule cells.**

(**A**) Simultaneous recording of GCaMP8s signals and cell-attached recording of APs in a granule cell illustrated schematically. Granule cell responses were evoked by electrically stimulating the medial perforant path.

(**B**) Sub- (top) and suprathreshold (bottom) GCaMP8s signals in a granule cell evoked by stimulating the medial perforant path. The pipette for the simultaneous cell-attached recording was depicted schematically.

(**C**) GCaMP8s signals in the granule cell in **B** plotted against time. The intensity of the electrical stimulation was varied to evoke a mixture of sub- and suprathreshold responses, which were classified using simultaneously recorded electrophysiological signals. Arrows indicate the timing of the synaptic stimulation.

(**D**) Cell-attached recordings corresponding to the sub- and suprathreshold $Ca^{2+}$ signals from the responses highlighted in blue and green in **C**.

(**E**) Summary graph comparing the peak amplitude of subthreshold $Ca^{2+}$ signals (blue) with that of suprathreshold $Ca^{2+}$ signals (green). * indicates that $P < 0.001$ (two-sided Wilcoxon rank sum test). Filled circles indicate individual data points. Whiskers in box charts indicate the 2.5th and 97.5th percentiles of these data points, and the box itself indicates the mean, first quartile, and third quartile of the data points. 0.24 $\Delta F/F_0$ (dashed line) corresponds to three standard deviations above the mean of subthreshold $Ca^{2+}$ signals, which were identified by the absence of APs in the concurrent electrophysiological signals. This value, which separated supra- from subthreshold $Ca^{2+}$ signals, was used to identify granule cells activated by synaptic inputs in **Figs. 2G**–**I** and **8G–K**.

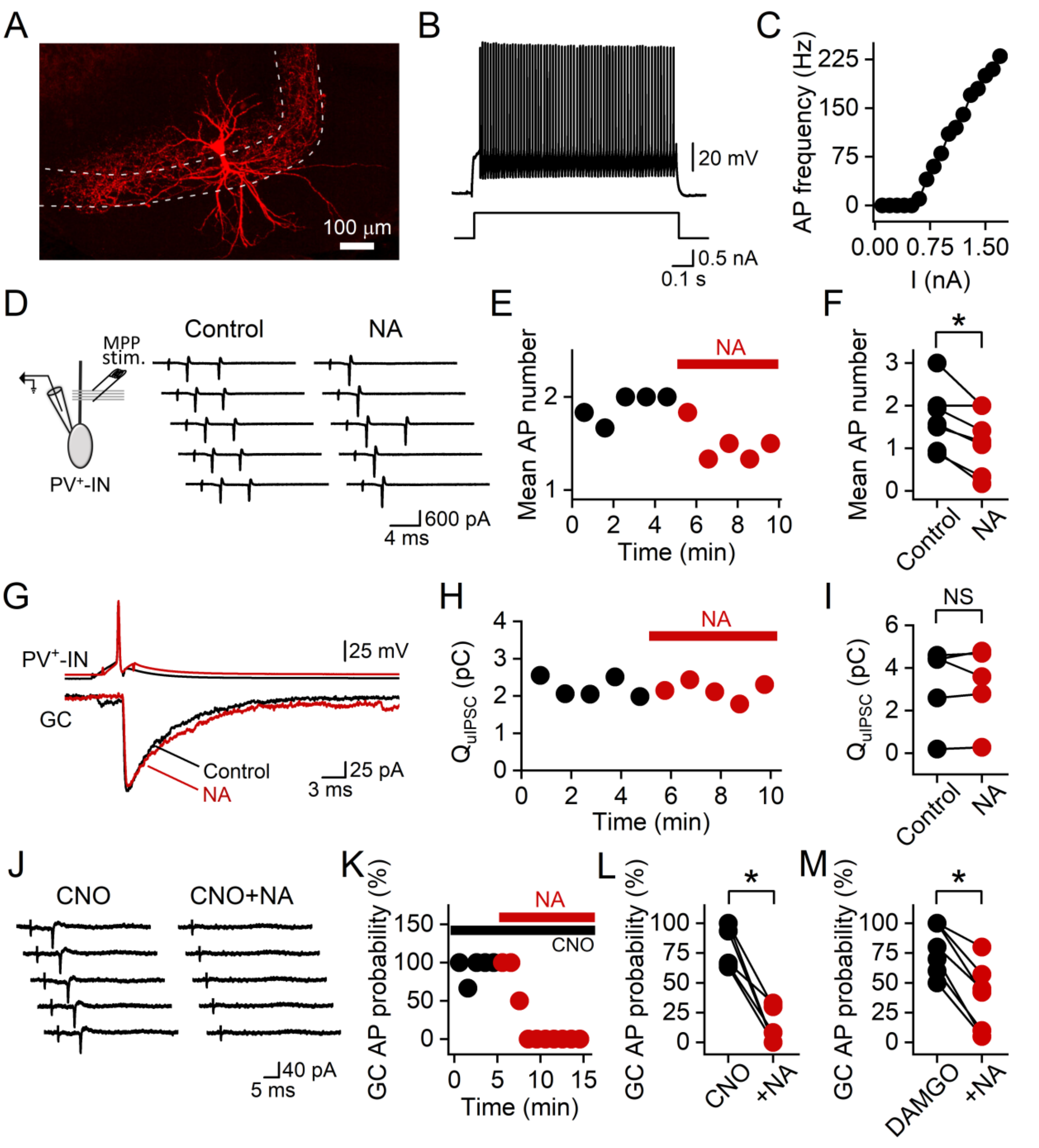


**Supplementary Figure 2. The inhibition of granule cells by noradrenaline is independent of PV$^+$-INs.**

(**A**) Morphology of a PV$^+$-IN in the dentate gyrus. White dashed lines indicate the borders of the granule cell layer.

(**B**) APs (top) in a PV$^+$-IN evoked by somatic injection of a 1-s depolarizing current pulse (bottom).

(**C**) Steady-state AP frequency plotted against the amplitude of the depolarizing current pulses from the experiment in **B**.

(**D**) Cell-attached recording of APs in a $PV^+$-IN, evoked by medial perforant path inputs before and during the bath application of noradrenaline. Current traces from five successive trials were plotted for each condition.

(**E**) Mean number of $PV^+$-IN APs evoked by the synaptic stimulation plotted against time from the experiment in **D**. Each data point represents the mean value of six consecutive trials.

(**F**) Summary graph showing that noradrenaline reduced the mean number of APs in $PV^+$-INs evoked by medial perforant path inputs. * indicates that P = 0.03 (n = 7 cells, two-sided Wilcoxon signed-rank test).

(**G**) Top, APs in a $PV^+$-IN under control conditions (black) and in noradrenaline (red). Bottom, corresponding uIPSCs in a granule cell.

(**H**) uIPSC charge plotted against time from the experiment in **G**.

(**I**) Summary graph comparing uIPSC charge under control conditions with that in noradrenaline. NS indicates P = 0.5 (n = 5 experiments, two-sided Wilcoxon signed-rank test).

(**J**) Cell-attached recording of APs in a granule cell evoked by stimulating medial perforant path axons in a hippocampal slice, following viral expression of hM4Di in $PV^+$-INs. CNO (1 μM) was present in the extracellular solution throughout the experiment to inhibit $PV^+$-INs. Note that chemogenetic inhibition of $PV^+$-INs did not prevent the suppression of granule cell activity by noradrenaline.

(**K**) The probability for the synaptic stimulation to evoke an AP in the granule cell plotted against time from the experiment in **J**.

(**L**) Summary graph showing that chemogenetic inhibition of $PV^+$-INs did not prevent the inhibition of granule cells by noradrenaline. * indicates that P = 0.02 (n = 7 cells, two-sided Wilcoxon signed-rank test).

(**M**) Similar to **L**, but the experiments were performed in the presence of bath-applied DAMGO to inhibit $PV^+$-basket cells throughout the experiment. * indicates that P = 0.02 (n = 7 cells, two-sided Wilcoxon signed-rank test).

In **F**, **I**, **L**, and **M**, data points from the same experiment are connected by a line.

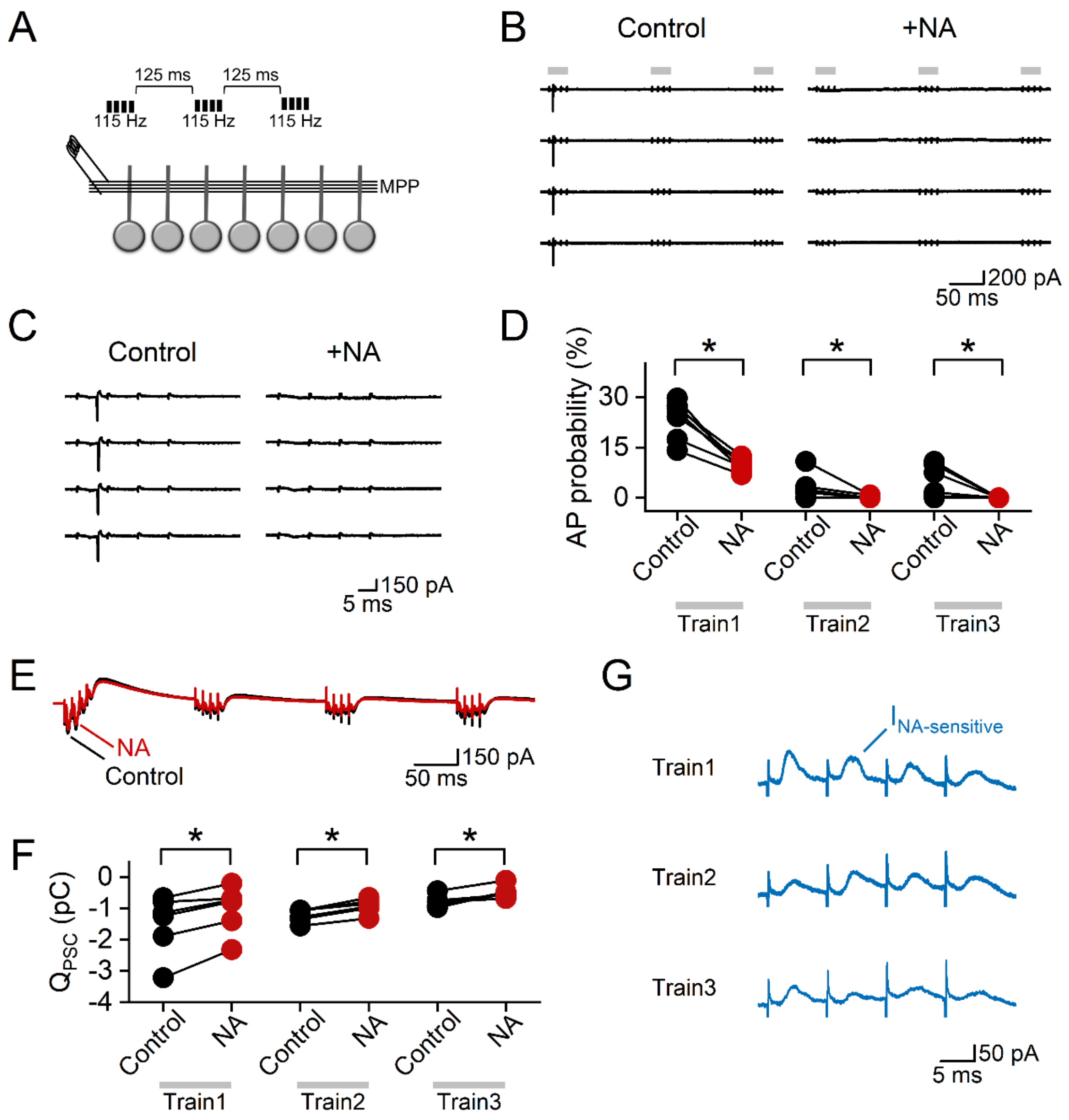


**Supplementary Figure 3. Noradrenaline inhibits granule cell firing evoked by repetitive stimulation of medial perforant path axons.**

(**A**) Schematic of the experimental design, showing three short trains of 115-Hz electrical stimulation delivered to medial perforant path axons, with a 125-ms interval between successive trains.

(**B**) Cell-attached recording from a granule cell showing that noradrenaline inhibited APs evoked by this stimulation protocol. Grey bars indicate the time windows for delivering the synaptic stimuli.

(**C**) From the same experiment in **B**, showing the response of the cell to the first train of the synaptic stimulation on expanded scales.

(**D**) Summary graph showing the probability of each train to activate the granule cell under control conditions and in noradrenaline.

(**E**) Superimposition of postsynaptic currents evoked by the stimulation protocol under control conditions and in noradrenaline.

(**F**) Charge of postsynaptic currents evoked by the first, second, and third stimulation trains under control conditions and in noradrenaline.

(**G**) Noradrenaline-sensitive postsynaptic currents evoked by the first, second, and third stimulation trains.

In summary plots, data points from the same experiment are connected by a line.

* indicates that P = 0.01–0.04 (n = 6 experiments, two-sided Wilcoxon signed-rank test).

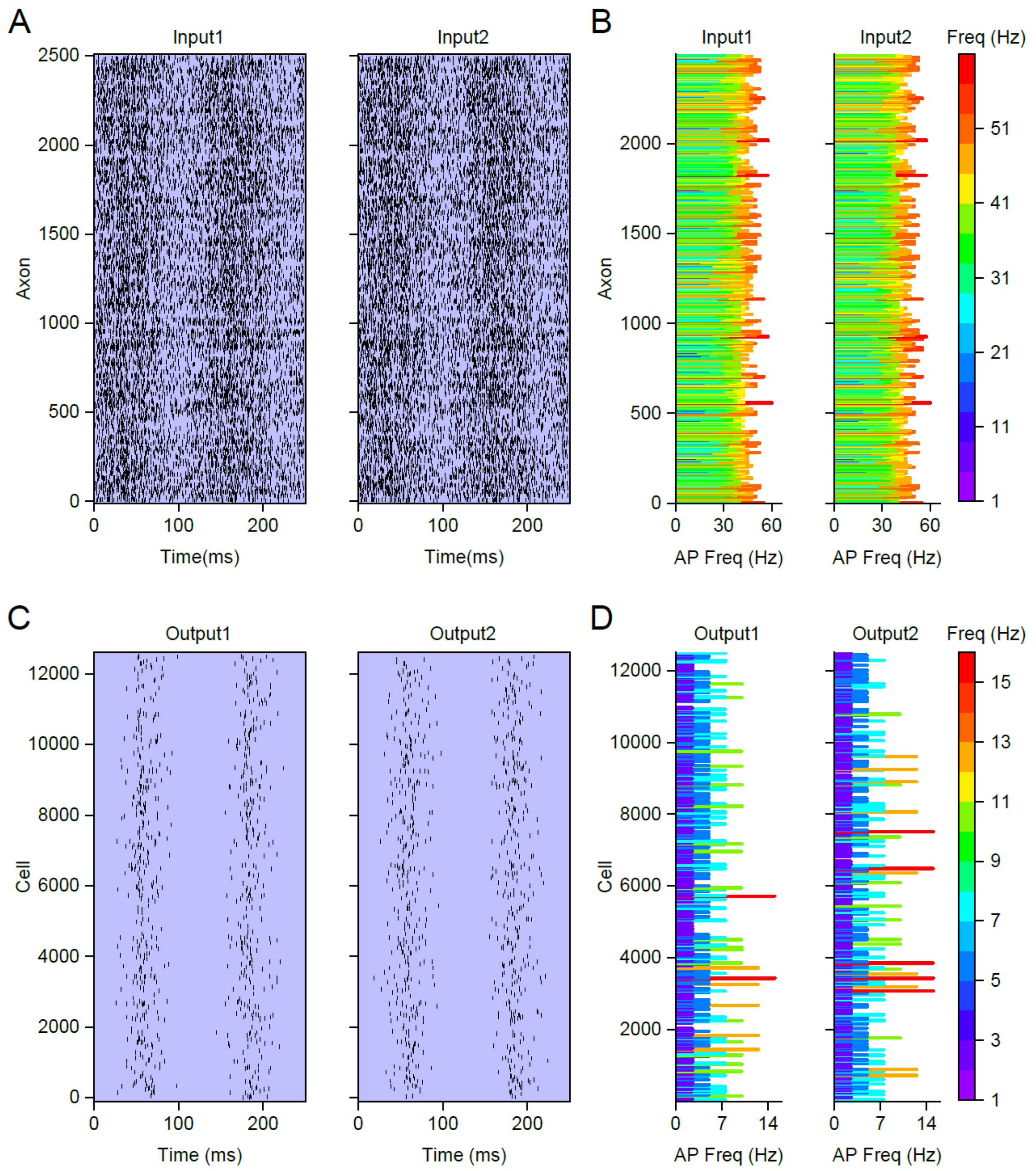


**Supplementary Figure 4. A simulation of decorrelation performed by feedforward inhibition.**

(**A**) Raster plots showing a pair of nearly identical input signals (Input1 and Input2, correlation = 0.82) in 2500 presynaptic axons. Each symbol represents a spike in an axon. The timing of axonal APs was determined by a random Poisson process and modulated by an 8-Hz sinusoidal function to mimic theta modulation of excitatory inputs from the entorhinal cortex[1].

(**B**) Axonal population vectors derived from raster plots in **A**, representing the steady-state AP rate in each axon. For visual comparison, AP frequency in each axon was also represented by a color code.

(**C**–**D**) Similar to **A**–**B**, but showing the output signals generated by 12500 postsynaptic excitatory neurons (Output1 and Output2, correlation = 0.49) in response to the two input signals in **A**.

To reduce the computational load of the simulation, the presynaptic network included 2500 axons. Individual neurons in the postsynaptic network sampled inputs from 2% of the presynaptic axons[2], corresponding to 50 randomly selected axons. This degree of convergence is relatively sparse in comparison to the innervation of individual granule cells by perforant path axons in the rodent brain[2]. To compensate for the low excitatory drive generated by the sparse connectivity in the model, the mean AP frequency in active presynaptic axons (36 Hz) was an order of magnitude higher than the mean spike rate in layer 2 neurons of the rodent entorhinal cortex during spatial exploration[3].

**Supplementary Table 1. Titer of AAVs**

| | Addgene catalog number | Titer |
|---|---|---|
| pAAV-Syn-FLEX-rc[ChrimsonR-tdTomato] | 62723 | $5 \times 10^{12}$ vg $mL^{-1}$ |
| pGP-AAV-syn-jGCaMP8s-WPRE | 162374 | $1 \times 10^{13}$ vg $mL^{-1}$ |
| pAAV-mDlx-NLS-mRuby2 | 99130 | $1 \times 10^{11}$ vg $mL^{-1}$ |
| pAAV-hSyn-DIO-hM4D(Gi)-mCherry | 44362 | $7 \times 10^{12}$ vg $mL^{-1}$ |
| pAAV-hDlx-Flex-GFP | 83895 | $7 \times 10^{12}$ vg $mL^{-1}$ |

**Supplementary Table 2. Coordinates of stereotaxic viral injections**

| Region | Coordinates from bregma | | |
|---|---|---|---|
| | anteroposterior (AP) | mediolateral (ML) | dorsoventral (DV) |
| Locus coeruleus (500 nL per site) | –5.4 mm | ± 0.8 mm | –3.4 mm |
| Hippocampus (100 nL per site) | –1.95 mm | ± 1.4 mm | –2.1 mm, –1.9 mm |
| | –3.6 mm | ± 2.5 mm | –3 mm |

**Supplementary Table 3. Passive and active properties of excitatory and inhibitory interneurons in the simulations**

| | | Excitatory neurons | Inhibitory interneurons |
|---|---|---|---|
| Morphology | | A spherical soma connected to a 210-μm-long apical dendrite | A spherical soma |
| Passive properties | | | |
| Leak conductance (A) | Density | 0.03 mS $cm^{-2}$ in the soma and dendrite | 0.05 mS $cm^{-2}$ in the soma |
| | Reversal potential | –80 mV | –62 mV under control conditions or –51 mV to mimic the modulation by noradrenaline |
| Membrane capacitance | | 1 μF $cm^{-2}$ | 1 μF $cm^{-2}$ |
| Cytoplasmic resistivity (B) | | 194 Ω cm | 100 Ω cm |
| Active properties | | | |
| Voltage-gated $Na^+$ conductance (C) | Density | 0.1 and 0 S $cm^{-2}$ in the soma and dendrite, respectively | 0.03 S $cm^{-2}$ in the soma |
| Voltage-gated $K^+$ conductance | Density | 0.075 S $cm^{-2}$ in the soma and dendrite | 0.015 S $cm^{-2}$ in the soma |
| Description of the voltage-gated $Na^+$ conductance ($g_{Na}$) (D) | | $g_{Na} = \overline{g_{Na}} \times m^3 \times h \times s$ where $\frac{dm}{dt} = \frac{m_\infty - m}{\tau_m}$, $m_\infty = \frac{\alpha_m}{\alpha_m + \beta_m}$, and $\tau_m = \frac{1}{(\alpha_m + \beta_m) \times 2.46}$ | |

| | | |
|---|---|---|
| | $\frac{dh}{dt} = \frac{h_\infty - h}{\tau_h}$, and $\tau_h = \frac{1}{(\alpha_h + \beta_h) \times 2.46}$ $\frac{ds}{dt} = \frac{s_\infty - s}{\tau_s}$ | |
| $\alpha_m$ | $\frac{0.4 \times (V + 23.7)}{1 - e^{\frac{V + 23.7}{-7.2}}}$ | $\frac{0.4 \times (V + 20)}{1 - e^{\frac{V + 20}{-7.2}}}$ |
| $\beta_m$ | $\frac{-0.124 \times (V + 23.7)}{1 - e^{\frac{V + 23.7}{7.2}}}$ | $\frac{-0.124 \times (V + 20)}{1 - e^{\frac{V + 20}{7.2}}}$ |
| $h_\infty$ | $\frac{1}{1 + e^{\frac{v + 48.7}{4}}}$ | $\frac{1}{1 + e^{\frac{v + 50}{4}}}$ |
| $\alpha_h$ | $\frac{0.03 \times (V + 43.7)}{1 - e^{\frac{V + 43.7}{-1.5}}}$ | $\frac{0.03 \times (V + 45)}{1 - e^{\frac{V + 45}{-1.5}}}$ |
| $\beta_h$ | $\frac{-0.01 \times (V + 43.7)}{1 - e^{\frac{V + 43.7}{1.5}}}$ | $\frac{-0.01 \times (V + 45)}{1 - e^{\frac{V + 45}{1.5}}}$ |
| $s_\infty$ | 1 | 1 |
| $\tau_s$ | $\frac{e^{0.09 \times (V + 58.7)}}{0.0003 \times [1 + e^{0.45 \times (V + 58.7)}]}$ | $\frac{e^{0.09 \times (V + 60)}}{0.0003 \times [1 + e^{0.45 \times (V + 60)}]}$ |
| Description of the voltage-gated K+ conductance ($g_K$) (D) | $g_K = \overline{g_K} \times n$ where $\frac{dn}{dt} = \frac{n_\infty - n}{\tau_n}$ | |
| $n_\infty$ | $\frac{1}{1 + e^{0.112 \times (19.5 - V)}}$ | $\frac{1}{1 + e^{0.112 \times (19.5 - V)}}$ |
| $\tau_n$ | $\frac{e^{0.079 \times (19.5 - V)}}{0.02 \times [1 + e^{0.112 \times (19.5 - V)}]}$ | $\frac{e^{0.079 \times (19.5 - V)}}{0.02 \times [1 + e^{0.112 \times (19.5 - V)}]}$ |

(**A**) Based on the density of the leak conductance, the membrane time constant was 31 ms for excitatory neurons and 20 ms for interneurons, respectively. Resting membrane potential and membrane time constant values of the excitatory neurons were based on Supplementary Reference 4[4], whereas those for interneurons matched the resting membrane potential and the membrane time constant of dentate gyrus $CCK^+$-INs in our experiments.

(**B**) The cytoplasmic resistivity of the excitatory neurons was based on Reference 5[5]. Hippocampal interneuron cytoplasmic resistivity values reported by previous studies vary by an order of magnitude[6, 7]. To minimize the impact of the cytoplasmic resistivity value on the voltage transfer in individual interneurons, interneurons in our model are represented by single-compartment isopotential units.

(**C**) To capture the strong attenuation of backpropagating APs in granule cell dendrites[8], excitatory neurons in our simulations do not express any dendritic voltage-gated $Na^+$ conductance.

(**D**) The voltage-gated $Na^+$ and $K^+$ conductance models were based on a modification of the models in Supplementary Reference 9[9].

**Supplementary Table 4. Summary of synaptic conductances in the simulations**

| | | Excitatory neurons | Inhibitory interneurons |
|---|---|---|---|
| Unitary AMPA conductance (A) | Peak amplitude | 19.2 pS | 15.3 pS |
| | Rise time constant | 1 ms | 1 ms |
| | Decay time constant | 5 ms | 3 ms |
| | Reversal potential | 0 mV | 0 mV |
| Unitary NMDA conductance (B) and (C) | Peak amplitude | 21.1 pS | 16.9 pS |
| | Rise time constant | 3 ms | 3 ms |
| | Decay time constant | 70 ms | 70 ms |
| | Reversal potential | 0 mV | 0 mV |
| Unitary GABA conductance (D and E) | Peak amplitude | 90 pS | |
| | Rise time constant | 1 ms | |
| | Decay time constant | 8.13 ms | |
| | Reversal potential | –75 mV | |

(**A**) Based on the peak amplitude of the unitary AMPA conductance, activating a single excitatory synapse in excitatory neuron dendrites generates a somatic EPSP with an amplitude of approximately 0.8 mV[8]. The peak amplitude of the unitary AMPA conductance in inhibitory neurons was adjusted so that the peak amplitude of somatic EPSCs in interneurons matched that of the excitatory neurons[10].

(**B**) The NMDA to AMPA conductance ratio in excitatory and inhibitory neurons was based on Supplementary References 11 and 12[11, 12].

(**C**) To capture its voltage-dependence, NMDA currents ($I_{NMDA}$) were simulated as

$$I_{NMDA} = g_{NMDA} \times (V - E_{NMDA}) / [1 + 0.3 \times \exp(-0.08 \times V)]$$

where V is the membrane potential, $g_{NMDA}$ is the NMDA conductance, and $E_{NMDA}$ is the reversal potential of NMDA currents[13].

(**D**) The peak amplitude of unitary GABA conductance was calibrated so that activation of a single inhibitory synapse produced an IPSP of approximately 1 mV from a membrane potential of –60 mV at the soma.

(**E**) The decay time constant of the unitary GABA conductance was based on our recordings from monosynaptically connected $CCK^{+}$-IN and granule cell pairs (8.13 ± 1.6 ms, n = 6 experiments).
